# Nutritional supplementation with vitamin E or plant extracts affects redox and immune response in early lactating dairy cows

**DOI:** 10.1101/2025.09.19.677329

**Authors:** Angélique Corset, Anne Boudon, Aude Remot, Benoît Graulet, Jean-François Ricouleau, Sabrina Philau, Colette Mustiere, Laurence Le-Normand, Ophélie Dhumez, Pierre Germon, Marion Boutinaud

## Abstract

In dairy cows, early lactation is a period prone to oxidative stress, inflammation and health problems. Our objective was to investigate the effects of the nutritional supplementation of early lactating cows with plant extracts or vitamin E on physiological and immunological status. Forty-five Holstein cows were divided into three groups: control group (n = 15), vitamin E group (n = 16) (3,000 IU/d for 3 weeks before and 1,000 IU/d after calving), and plant extract group (n = 14) (10 g/d only after calving). Their redox and immune status were monitored during the first 12 weeks of lactation, every four week. In week 12, plasma malondialdehyde levels in plasma were lower in the vitamin E and plant extract groups than in the control group. The vitamin E and plant extract groups had lower *ex vivo* plasma CCL4, IL6 and IL8 cytokine levels, lower proportions of blood regulatory neutrophils, and less ROS production by blood neutrophils than the control group. In terms of immune response, vitamin E and plant extracts down-regulated the gene expression of *FABP3* in the milk mammary epithelial cells of multiparous cows. Therefore, supplementation with either vitamin E or plant extracts could prevent systemic hyper-inflammation and improve the immune response during early lactation.

## INTRODUCTION

The onset of lactation in dairy cows induces dramatic changes in metabolic fluxes, driven by mammary gland demands that challenge many homeostatic regulations (1). These changes, and particularly those related to energy metabolites, are often associated with systemic inflammation and oxidative stress (2, 3). Altered immunity during early lactation has also been reported, so that when cows are exposed to pathogens, there is a greater risk of them moving from a healthy state to an infectious pathological state (4, 5), mastitis being the most common disease. Mastitis is associated with reduced milk production and an increase in the number of milk cells and it also can result in tissue damage in the mammary gland (6). Moreover, mammary epithelium integrity also appears to be affected by oxidative stress and inflammation (7). More specifically, disruption of the integrity of this epithelium can induce cell apoptosis and has been seen to reduce synthetic activity in lactating goat mammary glands (8).

The use of vitamin E supplementation in dairy cows has long been known to limit the incidence of mastitis (9). For instance, peripartum vitamin E supplementation has been shown to reduce oxidative stress, improve neutrophil migration, lower somatic cell counts in milk and reduce the number of mastitis cases (10–12). The NASEM recommends vitamin E supplementation of 3000 IU/d during the last 3 weeks of gestation and 1000 IU/d for lactating dairy cows (13). Plant extracts can be added to the diet to improve the antioxidant and immune functions, a strategy that has already been proposed for dairy cows or other farm animals. *Silybum marianum* induced an improvement in the immune function (i.e. increased phagocytosis of monocytes) and an increase in milk production, while preventing liver damage in dairy cows (14, 15). The use of *Arctium lappa* showed a reduction in oxidative stress in rats (i.e. a reduction in lipid peroxidation) (16). In the rabbit heart and kidney, *Salix alba* induced a reduction in lipid oxidation, while the activities of antioxidant enzymes (catalase, glutathione peroxidase (GPx)) were improved (17). However, studies to test the efficiency of plant supplementation in dairy cows remain scarce and only a few studies have so far explored the benefits of combining the properties of several plant extracts in a supplementation strategy. Using a product that contains of a combination of plants could jointly improve the redox and immune status of cows.

The objective of this study was therefore to evaluate the effects on the redox and immune status and mammary tissue damage in dairy cows in early lactation of two nutritional strategies during the first 12 weeks of lactation, based on supplementation with either vitamin E or a combination of plant extracts, compared with an unsupplemented nutritional strategy (‘control’). Supplementation with vitamin E was considered to be a positive control because the effects of vitamin E on the inflammatory response in early-lactation cows have been well described (18). Based on previous findings (19), we hypothesized that supplementation with vitamin E or plant extracts would improve the antioxidant response and reduce inflammation, and might have an effect on mammary epithelium integrity.

## RESULTS

### Plasma and ingested vitamin E, and plasma metabolites

Milk yield and dry matter intake (DMI) increased in line with parity and week of lactation (*P* < 0.001, Fig 1a and 1b) but were not affected by the treatment. Considering the vitamin E content of the die, the supplementation, and individual DMI, we showed that cows in the vitamin E group ingested more α-tocopherol than control and plant extract cows (*P* < 0.001, Fig 1c). Ingested γ-tocopherol increased in line with parity and levels were higher during lactation than during late gestation in all three groups (*P* < 0.001, Fig 1d).

**Figure 1:**
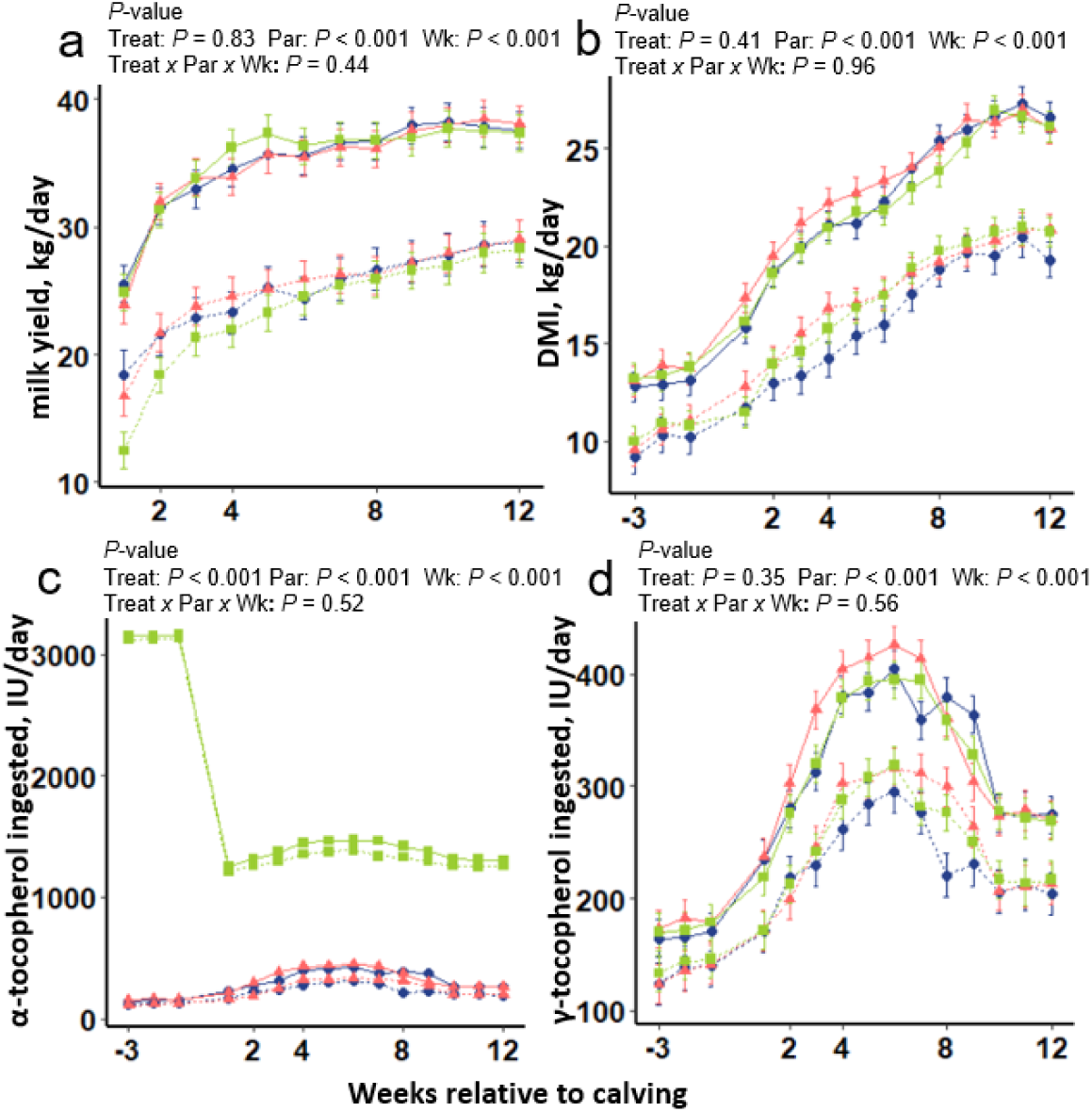
Milk yield in kg/d, dry matter intake (DMI) in kg/d, and ingested amounts of vitamin E (α and γ) in IU/d in control (n = 15), vitamin E (n = 16), and plant extract (n = 14) groups of dairy cows (primiparous and multiparous cows) from 3 weeks before calving to 12 weeks of lactation. Primiparous cows shown as dotted lines (---) and multiparous as solid lines (□). The control group is represented by solid and dotted red lines (▲), the vitamin E group by solid and dotted green lines (■), and the plant extract group by solid and dotted blue lines (●). Adjusted means and SEM are represented and data were analyzed according to a mixed model. Significant differences between parity (Par), treatment (Treat) and week (Wk) and their interaction (Treat x Par x Wk) are noted above the graphs.

Plasma urea, glucose, non-esterified fatty acids (NEFA), β-hydroxybutyrate (BHB) and calcium concentrations displayed significant variations over time as expected but were not affected by vitamin E and plant extracts except for NEFA, which tended to be more elevated in the plant extract group than in the vitamin E group (*P* = 0.07) (Supplementary Table S1). NEFA and BHB levels were higher in weeks 2 and 4 than in weeks 8 and 12 (*P* < 0.001). Plasma urea and glucose levels were lower in weeks 2 and 4 than in weeks 8 and 12 (*P* < 0.001), while plasma calcium was lower on day 1 after calving than on day 3 and in weeks 2, 4, 8 and 12 (*P* < 0.001).

Plasma levels of vitamin E (α-tocopherol, γ-tocopherol) and vitamin A (trans-βcarotene, 9cis-βcarotene, 13cis- βcarotene) fell after calving, and then rose from week 4 to week 12 of lactation (*P* < 0.001), except for retinol (*P* = 0.10) (Fig. 2). Plasma α-tocopherol levels were higher in the vitamin E group than in the control group (4.74 ± 0.19 *vs* 3.95 ± 0.28 µg/ml, *P* = 0.04, Fig. 2a). Plasma γ-tocopherol levels were higher in the control and plant extract groups than in the vitamin E group (0.72 ± 0.02 and 0.73 ± 0.03 *vs* 0.64 ± 0.02 µg/ml, *P* = 0.01, Fig. 2b). Plasma trans-βcarotene levels tended to be higher in the plant extract group than in the vitamin E group (1.39 ± 0.08 *vs* 1.15 ± 0.07 µg/ml, *P* = 0.08, Fig. 2e).

**Figure 2:**
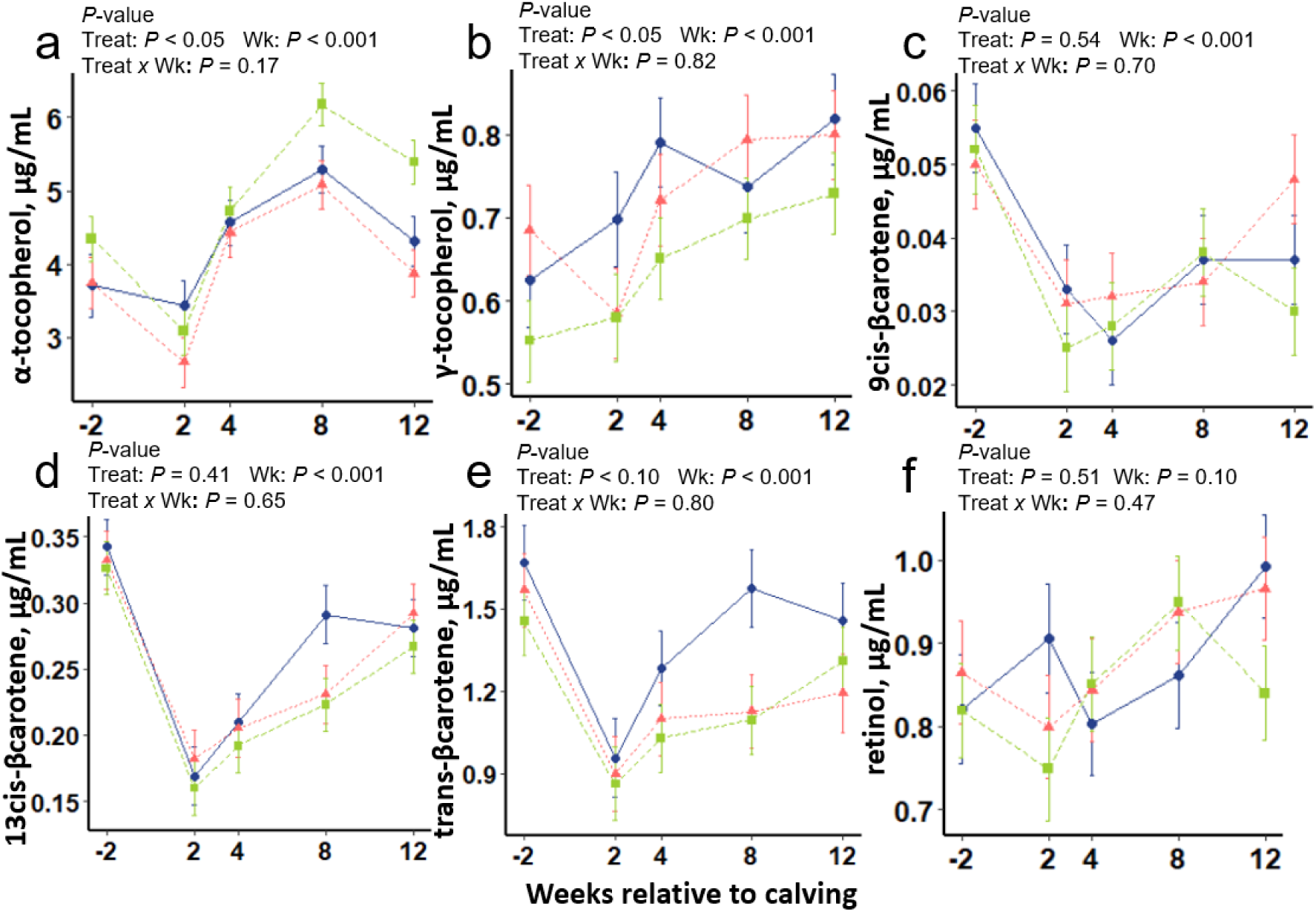
Plasma vitamin E and vitamin A levels in the control group (n = 15), vitamin E supplemented group (n = 16) and plant extract supplemented group (n = 14) of dairy cows in weeks -2, 2, 4, 8 and 12 relative to calving. Vitamin E can be present in plasma in several chemical forms: α-tocopherol, µg/mL (**A**), γ-tocopherol, µg/mL (**B**). Vitamin A can be present in plasma in several chemical forms: 9cis-βcarotene (**C**), 13cis-βcarotene (**D**), all trans-βcarotene (**E**), and retinol (**F**) in µg/mL. The control group is represented as a dotted red line (- -▲- -), the vitamin E group as a dotted green line (--■--) and the plant extract group as a solid blue line (□●□). Adjusted means and SEM are represented as a function of the week relative to calving, calculated according to a mixed model including a covariate measured in week -3. Significant differences between treatment (Treat) and week (Wk) and their interaction (Treat x Wk) are noted above the graphs.

### Oxidative stress and the antioxidant capacities of vitamin E and plant extracts in dairy cows

Plasma reactive oxygen metabolite (ROM) levels were higher in weeks 2, 4 and 12 than in week 8 of lactation (*P* < 0.001, Fig. 3a). Plasma ferric reducing antioxidant power (FRAP) was low in weeks 2, 4 and 8 and then rose in week 12 (*P* < 0.001, Fig. 3b). Plasma biological antioxidant potential (BAP) was higher in weeks 4 and 12 of lactation than in weeks 2 and 8 (*P* < 0.001, Fig. 3c) and the plasma oxidative stress indexes according to FRAP (OSI_FRAP_) or BAP (OSI_BAP_) were higher in weeks 2, 4 and 12 than in week 8 of lactation (*P* < 0.001) (Fig. 3d and 3e). Plasma GPx activity was higher in weeks 2, 4 and 12 than in week 8 of lactation (*P* < 0.001, Fig. 3f), while erythrocyte SOD activity was not impacted by week of lactation (*P* = 0.70, Fig. 3h). None of these variables (plasma ROM concentration, FRAP, BAP, OSI_FRAP_ and OSI_BAP_, GPx plasma activity, SOD erythrocyte activity) was affected by vitamin E and plant extracts (*P* > 0.10, Fig. 3). Erythrocyte GPx activity was lower in week 4 than in week 12 (*P* = 0.01) and the vitamin E group tended to exhibit more erythrocyte GPx activity than the plant extract group (4.36 ± 0.004 *vs* 4.35 ± 0.005 nmol/min/mL, *P* = 0.07) (Fig. 3g). Plasma SOD activity was low during the first week of lactation and then rose gradually (*P* < 0.01, Fig. 3h) in week 12. The vitamin E group tended to exhibit more plasma SOD activity than the plant extract group in week 12 (12.5 ± 0.86 *vs* 9.55 ± 0.95 nmol/min/mL, *P_treatment×week_*= 0.06) (Fig. 3h). Plasma malondialdehyde (MDA) levels were low between weeks 2 and 8, and then rose in week 12 of lactation (*P* = 0.003). Throughout the experiment, plasma MDA levels tended to be lower in the vitamin E group than in the control group (1.87 ± 0.15 *vs* 2.37 ± 0.16 nmol/mL, *P* = 0.09, Fig. 3j). Although the treatment×week interaction was not significant, simple comparisons of means by student’s tests showed evidences at week 12 that plasma MDA levels was lower in the vitamin E than in the control group (2.17 ± 0.21 *vs* 2.91 ± 0.23 nmol/mL, *P* = 0.04), and tended to be lower in the plant extract group than in the control group (2.18 ± 0.25 *vs* 2.91 ± 0.23 nmol/mL, *P* = 0.09) (Fig. 3j).

**Figure 3:**
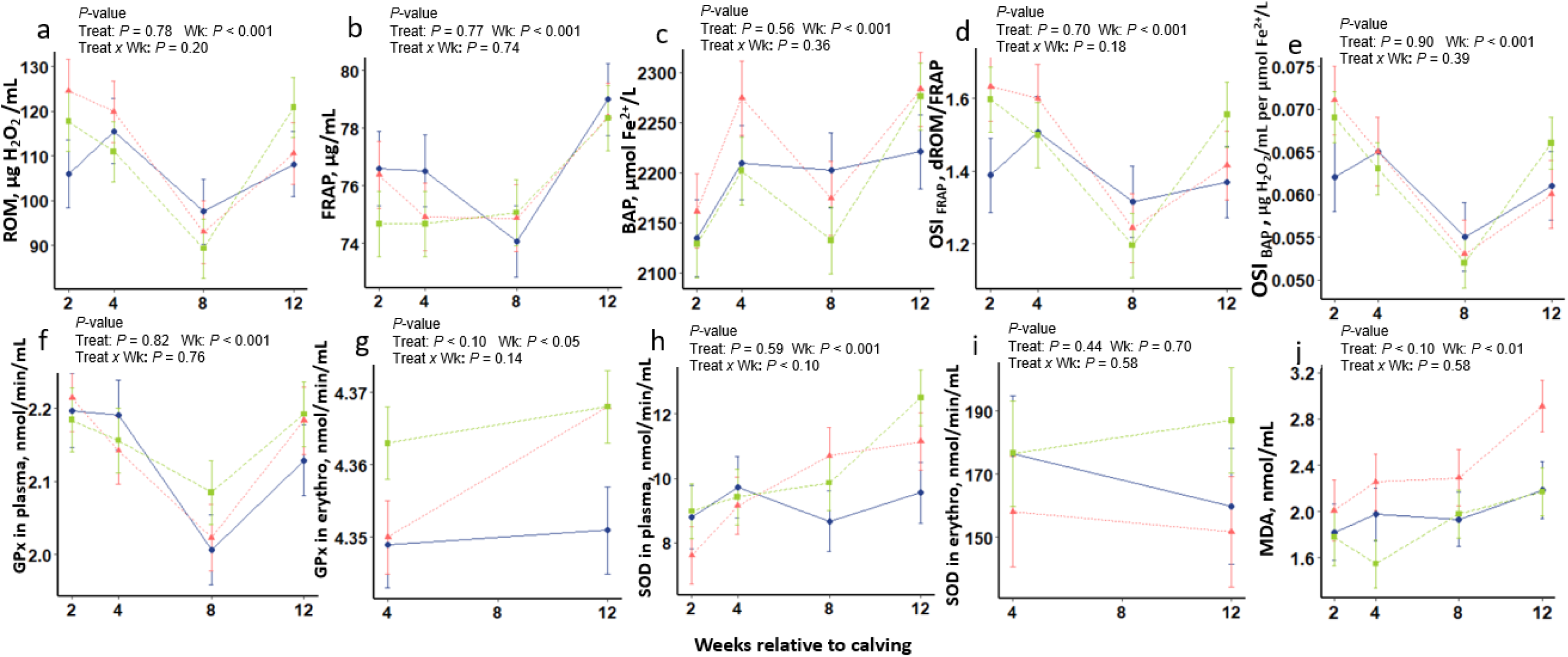
Biomarkers of redox balance and enzymatic activities in plasma or erythrocytes in the control group (n = 15), vitamin E group (n = 16), and plant extracts group (n = 14) of dairy cows at weeks 2, 4, 8 and 12 relative to calving. Reactive oxygen metabolite (ROM), µg H_2_O_2_ /mL (**A**), ferric reducing antioxidant power (FRAP), µg/mL (**B**), biological antioxidant power (BAP), µmol Fe^2+^/L (**C**), oxidative stress index OSI _FRAP_ (**D**), and OSI _BAP_, µg H_2_O_2_/mL per µmol Fe^2+^/L (**E**) were measured in plasma. The antioxidant enzyme activity of glutathione peroxidase (GPx) was measured in plasma (**F**) and erythrocytes (**G**). The antioxidant enzyme activity of superoxide dismutase (SOD) was measured in plasma (**H**) and erythrocytes (**I**). Plasma MDA (malondialdehyde) was represented in nmol/mL (**J**). The control group is represented as a dotted red line (- -▲- -), the vitamin E group as a dotted green line (--■--), and the plant extract group as a solid blue line (□●□). Adjusted means and SEM are represented as a function of the week relative to calving, calculated according to a mixed model including a covariate measured in week -3. Significant differences between treatment (Treat) and week (Wk) and their interaction (Treat x Wk) are noted above the graphs.

### Counting and measuring the immune capacities of immune cells

Plasma haptoglobin levels were higher in week 2 than in weeks 8 and 12 (*P* < 0.001, Fig. 4a). Plasma cortisol was lower in weeks 2 and 4 than in weeks 8 and 12 (*P* < 0.001, Fig. 4f). Plasma haptoglobin and cortisol levels were not affected by vitamin E and plant extracts (*P* > 0.10), and nor was the red blood cell count (P > 0.10, Supplementary Table S2). The white blood cell count was higher in week 2 than in weeks 4, 8 and 12 of lactation (*P* < 0.001, Fig. 4b). In multiparous cows, the white blood cell count tended to be lower in the plant extract group than in the control group (5.66 ± 0.66 *vs* 7.87 ± 0.63 10^9^/mL), while it was not affected by vitamin E and plant extracts in primiparous cows (treatment×parity*P_treatment × parity_* = 0.06, Fig. 4d). The monocyte count was higher in week 2 than in weeks 4, 8 and 12 of lactation (*P* < 0.001, Fig. 4c). In multiparous cows, the monocyte count was lower in the vitamin E group than in the control group (0.20 ± 0.04 *vs* 0.33 ± 0.04 10^9^/mL). Moreover, in primiparous cows, the plant extract group exhibited a higher blood monocyte count than the control group (*P_treatment×parity_* = 0.01, Fig. 4e). The blood neutrophil count was not affected by the week of lactation (*P* = 0.10, Fig. 4g). In the control group, the blood neutrophil count tended to be higher in multiparous than in primiparous cows (0.20 ± 0.04 *vs* 0.33 ± 0.04 10^9^/mL, *P* = 0.06), but this result was not observed in the vitamin E and plant extract groups (Fig. 4i).

**Figure 4:**
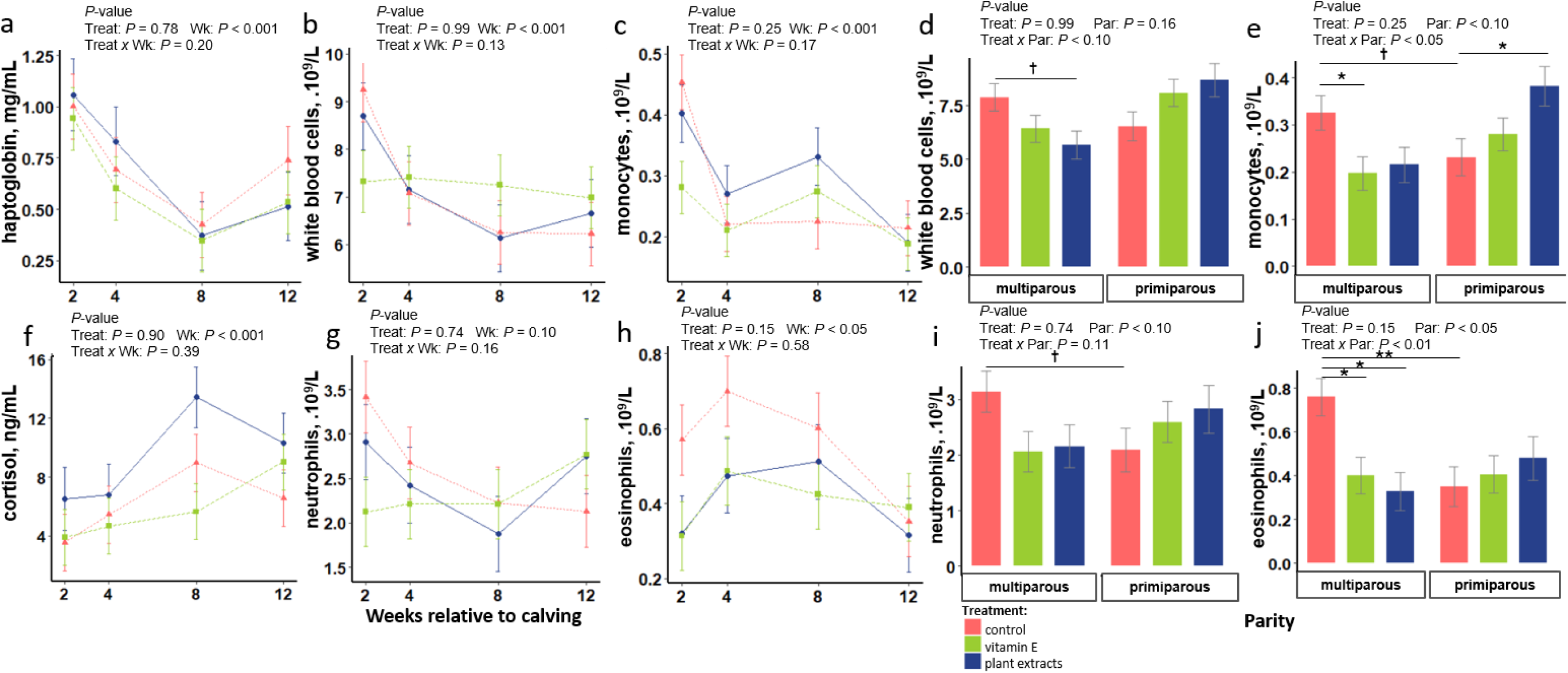
Plasma haptoglobin and cortisol concentrations and hematological profile in the control group (n = 15), vitamin E group (n = 16), and plant extracts group (n = 14) of dairy cows according to weeks 2, 4, 8 and 12 relative to calving, or according to parity in primiparous or multiparous cows. Haptoglobin, in mg/mL (**a**), white blood cells, in 10^9^/L, according to week relative to calving (**b**) or according to parity (**d**), monocytes, in 10^9^/L, according to week relative to calving (**c**) or according to parity (**e**), cortisol, in ng/mL (**f**), neutrophils, in 10^9^/L, according to week relative to calving (**g**) or according to parity (**i**), eosinophils, in 10^9^/L, according to week relative to calving (**h**) or according to parity (**j**). On the graph (**a, b, c, f, g, h**), the control group is represented as a dotted red line (- -▲- -), the vitamin E group as a dotted green line (--■--), and the plant extract group as a solid blue line (□●□). Adjusted means and SEM are represented as a function of the week relative to calving, calculated according to a mixed model including a covariate measured in week -3. Significant differences between treatment (Treat) and week (Wk) and their interaction (Treat x Wk), or parity (Par) and their interaction (Treat x Par) are noted above the graphs.

The blood eosinophil count was higher in weeks 4 and 8 than in week 12 of lactation (*P* = 0.02, Fig. 4h). In the control group of cows, the blood eosinophil count was higher in multiparous than in primiparous cows (0.76 ± 0.08 *vs* 0.35 ± 0.09 10^9^/mL, *P* = 0.01). In multiparous cows, the vitamin E and the plant extract groups exhibited higher eosinophil counts than the control group (0.40 ± 0.08 and 0.33 ± 0.09 *vs* 0.76 ± 0.08 10^9^/mL, *P_treatment×parity_* = 0.01, Fig. 4j).

The proportion of classical neutrophils, identified as negative for the class II major histocompatibility complex (MHCII-), was lower at week 4 than at weeks 8 and 12 (*P* < 0.001), and was higher in the vitamin E and plant extract groups than in the control group (95.73 ± 0.39 and 95.42 ± 0.44 *vs* 93.13 ± 0.49 %, *P* = 0.01) (Fig. 5a). Inverse variations were observed regarding the percentage of regulatory neutrophils expressing the class II major histocompatibility complex (*P* = 0.04, Fig. 5d).

**Figure 5:**
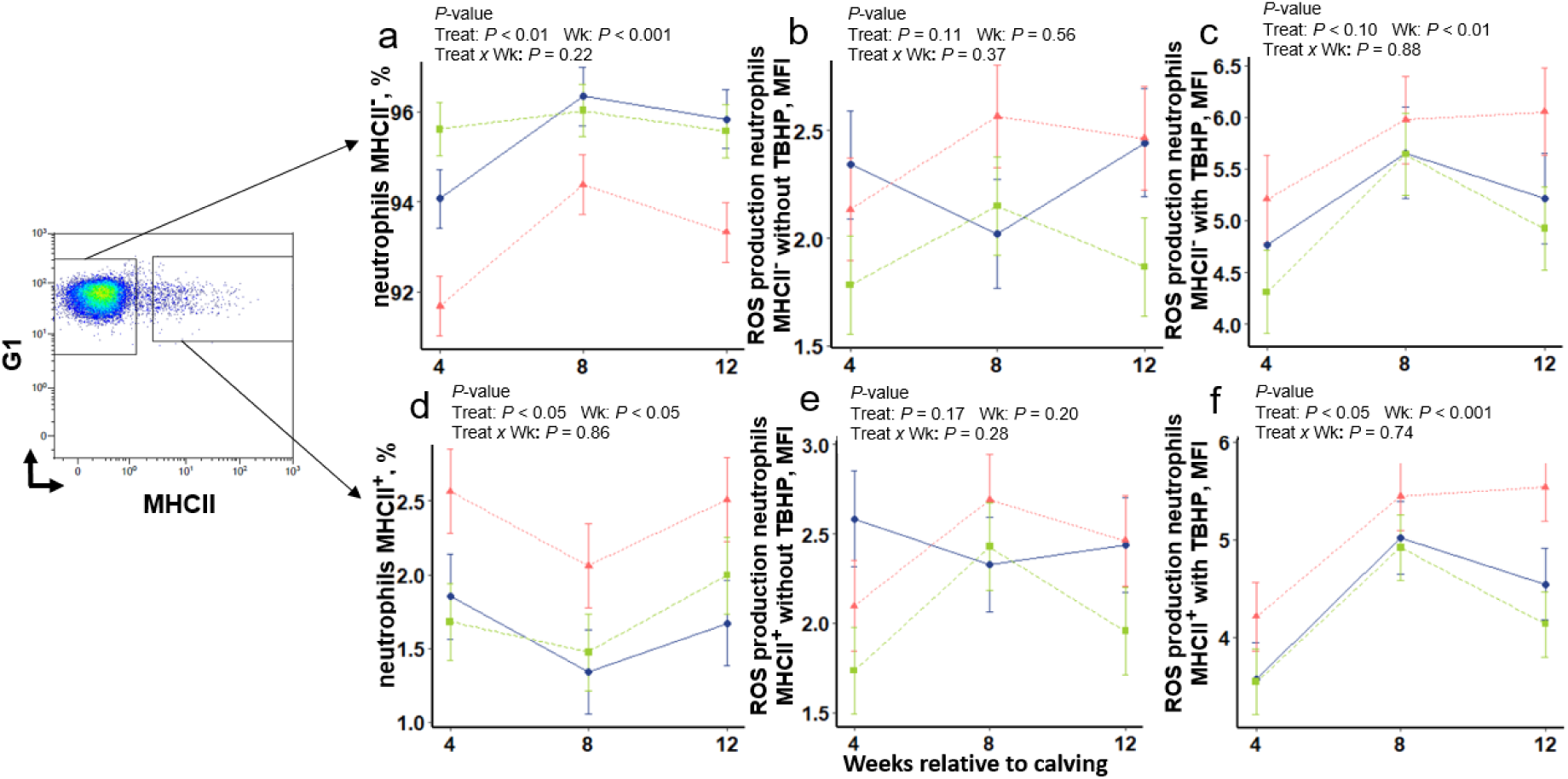
Flow cytometry identification of two types of neutrophils isolated from blood and measurement of their intracellular reactive oxygen species (ROS) production without or with stimulation *ex vivo* in the control (n = 15), vitamin E supplemented (n = 16), and plant extracts supplemented (n = 14) dairy cows at weeks 2, 4, 8 and 12 relative to calving. Percentages of neutrophils without class II major histocompatibility complex, MHCII**^−^** (**A**) and class II major histocompatibility complex neutrophils, MHCII**^+^** (**D**). Mean fluorescence intensity (MFI) measurement of intracellular ROS production by flow cytometry of MHCII**^−^** neutrophils without (**B**) or with (**C**) tert-butyl hydroperoxide (TBHP) stimulation. Measurement of intracellular ROS production of MHCII**^+^** neutrophils without (**E**) or with (**F**) TBHP stimulation. The control group is represented as a dotted red line (- -▲- -), the vitamin E group as a dotted green line (--■--), and the plant extract group as a solid blue line (□●□). Adjusted means and SEM are represented as a function of the week relative to calving, calculated according to a mixed model including a covariate measured in week -3. Significant differences between treatment (Treat) and week (Wk) and their interaction (Treat x Wk) are noted above the graphs.

The ability of animals to respond to infection was studied first by measuring the capacity of neutrophils to produce Reactive Oxygen Species (ROS) following stimulation with tert-butyl hydroperoxide (TBHP). Without TBHP stimulation, ROS production by classical and regulatory neutrophils was not affected by either vitamin E or plant extracts (*P* = 0.11 and *P* = 0.17, respectively) or week of lactation (*P* = 0.56 and *P* = 0.20, respectively) (Fig. 5b and 5e). After TBHP stimulation, classical neutrophils released higher levels of ROS at week 8 than at weeks 4 and 12 (*P* = 0.01, Fig. 5c), and regulatory neutrophils released higher levels of ROS at weeks 8 and 12 than at week 4 (*P* < 0.001, Fig. 5f). After TBHP stimulation, classical neutrophils tended to produce less ROS in the vitamin E and plant extract groups than in the control group (4.95 ± 0.23 and 5.21 ± 0.26 *vs* 5.74 ± 0.25 MFI (Mean Fluorescence Intensity), *P* = 0.07, Fig. 5c). Regulatory neutrophils had a lower ROS production in the vitamin E and plant extract groups than in the control group (4.20 ± 0.20 and 4.37 ± 0.22 *vs* 5.06 ± 0.21 MFI, *P* = 0.01, Fig. 5f).

The release of cytokines by blood cells after an ex vivo challenge with heat-killed (HK) *Escherichia coli* was higher in week 8 than in week 12 (*P* < 0.001, Fig. 6), particularly for CCL2, CCL4, CXCL10, IFNγ, IL1α, IL1β, IL2, IL6, IL8, TNFα, IL10, IL1Rα and IL4 (*P* < 0.05, Fig. 6). Whatever the week, the plasma mean log_10_ MFI (Median Fluorescence Intensity) values for CCL4 in the plant extract group and IL8 in the vitamin E and plant extract groups were lower than in the control group (*P* = 0.04 and *P* = 0.02, respectively, Fig. 6). The plasma log10 MFI for IL6 also tended to be lower in the vitamin E group than in the control group (*P* = 0.08, Fig. 6). For IL17A, statistical analysis highlighted a treatment×week interaction (*P =* 0.02). At week 8, the plant extract group had higher plasma log10 MFI values for IL17A than the control group (*P* = 0.03, Student’s t-test). At week 8, the plasma log10 MFI for IFNγ was also higher in the vitamin E and plant extract groups than in the control group (*P* = 0.06 and *P* = 0.10 respectively, Student’s t-test). Without the HK *E. coli* challenge, plasma cytokine production was not affected by vitamin E and plant extracts (*P* > 0.10, Supplementary Table S3).

**Figure 6:**
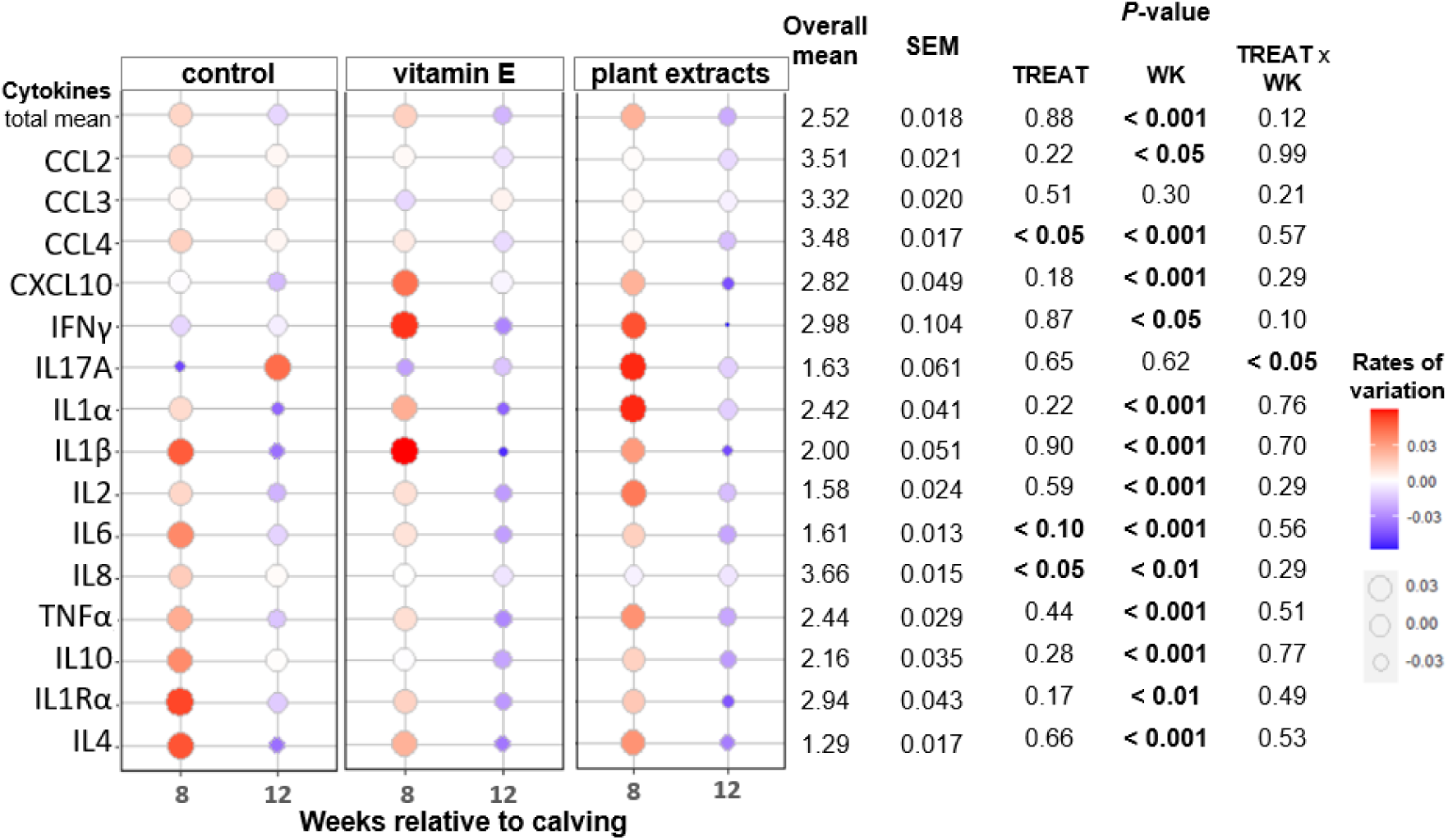
Measurement of 15 cytokines in plasma expressed in median fluorescence intensities (MFI) after an *ex vivo* heat killed *Escherichia coli* (HK *E. coli*) challenge in control (n = 15), vitamin E supplemented (n = 16), and plant extract supplemented (n = 14) dairy cows at weeks 8 and 12 relative to calving. The rates of variation for each cytokine was calculated to represent the difference between the control, vitamin E and plant extract groups according to the week of lactation. The total mean of cytokines (total mean), the chemokines (CCL2, CCL3, CCL4, CXCL10), pro-inflammatory cytokines (IFNγ, IL17A, IL1α, IL1β, IL2, IL6, IL8, TNFα) and anti-inflammatory cytokines (IL10, IL1Rα, IL4) were measured in plasma *ex vivo*. The cytokine quantities are represented with the rates of variation on the heat maps to observe the effect of treatment. The rate of variation was calculated according to the following formula: Rate of variation = (MFI mean - overall mean) / overall mean. The overall mean and standard error of the mean (SEM) were calculated for each cytokine. The probabilities were: TREAT = effect of parity, control vs. vitamin E vs. plant extracts; WK = effect of week, week 8 and 12 relative to calving; TREAT × WK = the interaction between parity and week. A covariate is chosen for the static calculations to highlight the effects of supplementation (vitamin E and plant extract groups).

### Mammary epithelium integrity and the expression of genes related to the immune response, and antioxidant and milk production functions in MEC

Levels of milk fat and protein and biomarkers of local mammary gland inflammation and mammary epithelium integrity were high at week 2 of lactation and then gradually decreased at weeks 4, 8 and 12 (somatic cell count (SCC), somatic cell score (SCS), mammary epithelial cell (MEC) concentration, MEC exfoliation into milk, Na^+^:K^+^ ratio in milk, Supplementary Table S4). None of these parameters was affected by vitamin E or plant extract supplementations (*P* > 0.10). Plasma lactose did not vary over treatment and week. Nevertheless, milk lactose levels were lower at weeks 4 and 8 than at week 12 (*P* = 0.01), and were lower in the vitamin E and plant extract groups than in the control group (*P* = 0.01), but these variations were not observed regarding the lactose yield (*P* = 0.65, Supplementary Table S4). Milk lactose concentrations were not affected by the interaction treatment by parity. However, simple comparisons of means by Student’s tests showed evidences that milk lactose concentrations were lower in the vitamin E and plant extract groups than in the control groups but only for multiparous cows (*P* < 0.001 and *P* = 0.01 for vitamin E and plant extracts, respectively).

The expression of several genes were evaluated in MEC isolated from milk (Supplementary Table S5). The mean expression of reference genes (*ACTB*, *GAPDH*, *PPIA*, and *RPLP0*) did not vary according to parity, treatment or week (*P* > 0.10, Fig. 7a, 7b, 7c). The expressions of antioxidant genes in milk MEC (namely *NFE2L2* and *GPX1)* were lower at week 4 than at week 12 (*P* < 0.001 and *P* < 0.001, Fig. 7a). At week 4, *NFE2L2* expression tended to be lower in the vitamin E group than in the plant extract group (-56 %, *P_treatment×week_* = 0.07, Fig. 7a), but *GPX1* expression did not vary significantly between groups (*P* = 0.94, Fig. 7a). The expressions of immune genes in milk MEC (namely *CXCL10*, *FABP3*, *IFN*β*1* and *LBP)* were higher at week 4 than at weeks 8 or 12 (*P* = 0.06, *P* < 0.001, *P* = 0.01, and *P* = 0.02 respectively, Fig. 7a). The *CXCL10* gene tended to be less strongly expressed in the vitamin E group than in the plant extract group (+ 99%, *P* = 0.07, Fig. 7a). As for the expression of the *FABP3*, *IFNB1* and *LBP* genes, we observed a significant effect of the treatment×parity×week interaction (Fig 7b). In multiparous cows at week 12, the *FABP3* gene was less strongly expressed in the vitamin E (-66%) and plant extract (-69%) groups than in the control group (*P_treatment×parity×week_*= 0.01, Fig. 7b). In week 4, the *IFN*β*1* gene was more strongly expressed in multiparous cows than in primiparous cows, and in the latter, it was also more strongly expressed in the vitamin E group than in the plant extract group (3.36-fold increase, *P_treatment×parity×week_*= 0.03, Fig. 7b). In multiparous cows at week 8, the *LBP* gene was less strongly expressed in the vitamin E group than the plant extract group (-81%). Moreover, in multiparous cows at week 12, the *LBP* gene was less strongly expressed in the plant extract group than in the control group (- 85%, *P_treatment×parity×week_* = 0.03, Fig. 7b). In primiparous cows, the *TJP1* gene tended to be more strongly expressed in vitamin E than in the control (1.45-fold increase) and plant extract groups (1.69-fold increase) (*P_treatment×parity_* = 0.07, Fig. 7c), while in the plant extract group, *TJP1* tended to be more strongly expressed in multiparous than in primiparous cows (1.19-fold increase), but parity did not affect the control group (*P* = 0.33). As an example of the milk synthesis genes influenced by week of lactation, *LALBA* tended to be less strongly expressed in week 4 than in week 8 (*P* = 0.06) but was not affected by vitamin E and plant extracts (*P* > 0.10, Fig. 7a).

**Figure 7:**
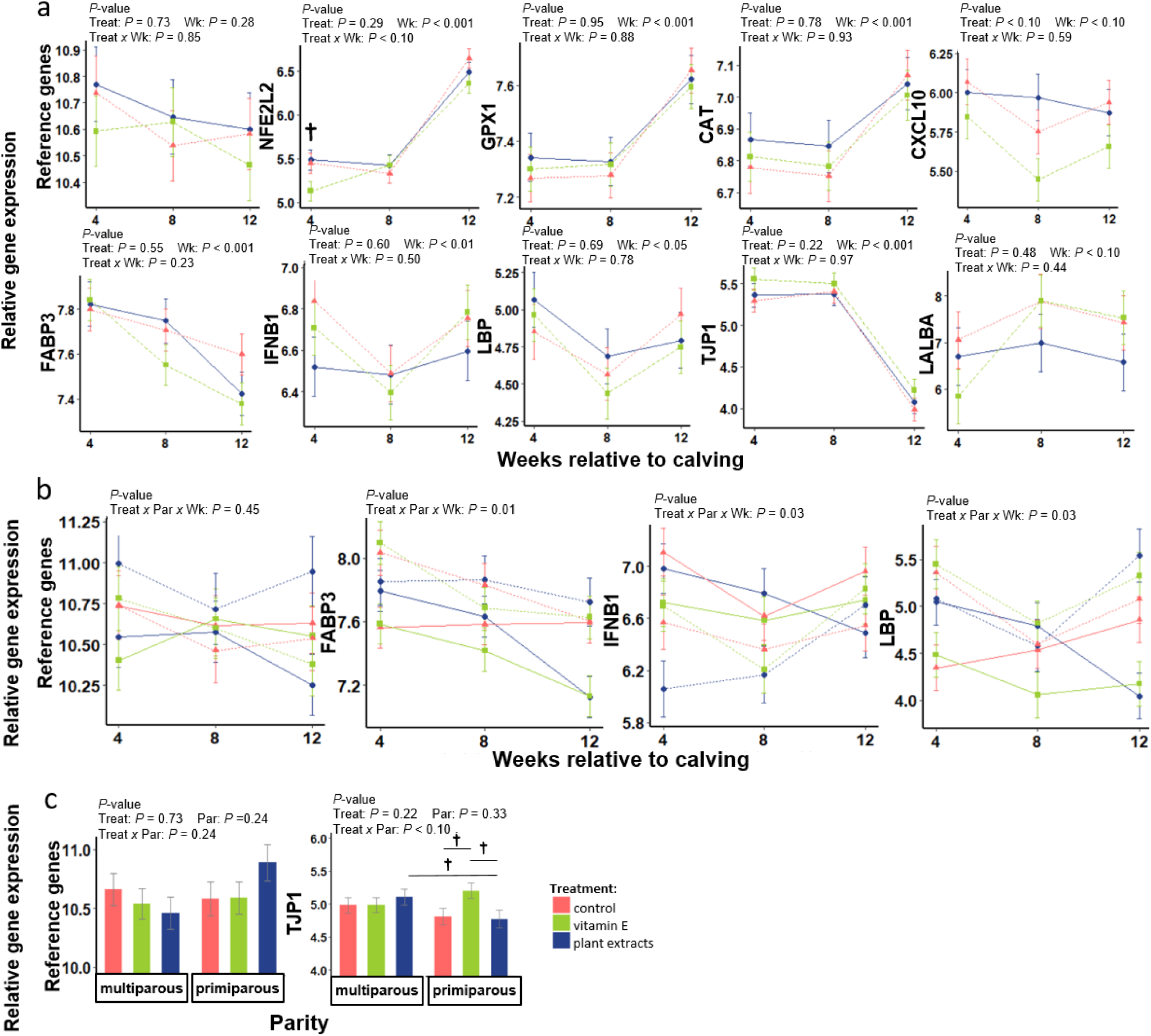
Abundance of antioxidant and immune function mRNA, and milk production mRNA determined by real-time quantitative PCR in milk isolated from mammary epithelial cells (MEC) in control (n = 15), vitamin E supplemented (n = 16), and plant extract supplemented (n = 14) dairy cows between weeks 4 and 12 relative to calving. On the graph (**a**), the control group is represented as a dotted red line (- -▲- -), the vitamin E group as a dotted green line (--■--), and the plant extract group as a solid blue line (□●□). On the graph (**c**), a triple interaction is represented with primiparous cows as a dotted line (---) and multiparous cows as a solid line (□) according to the control group as a solid and dotted red line (▲), the vitamin E group as a solid and dotted green line (■), and the plant extract group as a solid and dotted blue line (●). Adjusted means and SEM are represented as a function of the week relative to calving, calculated according to a mixed model. Significant differences between treatment (Treat) and week (Wk) and their interaction (Treat x Wk), or parity (Par) and their interaction (Treat x Par), and a triple interaction (Treat x Par x Wk) are noted above the graphs. Milk samples were collected on weeks 4, 8 et 12 after calving, then MEC from milk were prepared and an analysis of the gene expression involved in antioxidant, immune and milk production functions was made using *ACTB* (actin beta), *GAPDH* (Glyceraldehyde-3-Phosphate Dehydrogenase), *PPIA* (cyclophilin A), and *RPLP0* (Ribosomal Protein Lateral Stalk Subunit P0) as reference genes. The means and SEM (standard error of the mean) are represented.

## DISCUSSION

The objective of our study was to investigate the effects of supplementation with vitamin E and plant extracts on the redox and immune status and on mammary epithelial integrity in healthy dairy cows. None of the metabolite concentrations, whether NEFA and BHB for acetonemia, urea for uremia, glycemia for hypoglycemia and calcemia for milk fever, or the milk somatic cell count for mastitis, reached the clinically pathological alert thresholds (4, 20, 21). The feed analysis and intake measurements showed that findings in the vitamin E group were in line with our supplementation objectives, as confirmed by the presence of higher plasma α-tocopherol levels in the vitamin E group, and previously observed in other studies (11, 22). The higher plasma levels of all-trans-βcarotene in cows supplemented with plant extracts than those seen in the plasma of cows supplemented with vitamin E, indicated that these compounds were supplied by the plant extract.

The start of lactation in dairy cows is a period of oxidative stress where plasma ROM and OSI increase after calving, a finding in line with our observations (23). This is consistent with the high energy metabolism related to lactation between the time of calving and peak lactation. The absence of a vitamin E effect on plasma ROM in our study was consistent with the findings of another study (24). Our study confirmed a low level of antioxidant capacity in cows at the start of lactation, followed by an increase after the peak of milk yield, as previously observed (25). Our vitamin E supplementation did not increase total antioxidant capacity, unlike the results observed in another study involving supplementation with 3,000 IU for 8 weeks before calving (26). Few studies have examined these indicators (ROM, FRAP, BAP) in healthy cows under vitamin E supplementation (22, 24) and they did not reveal any effects of vitamin E supplementation at the same weeks as in our study.

The GPx antioxidant activity of plasma appeared to vary like ROM and OSI between weeks of lactation in response to oxidative stress, but without any effects of vitamin E and plant extracts. Previous studies had shown that plasma ROM and erythrocyte GPx activity were positively correlated (24). In our case, GPx activity in erythrocytes was higher in the vitamin E group when plasma ROMs were high, i.e. at week 4, in agreement with the effects of an intramuscular injection of vitamin E previously reported (27). Moreover, plasma SOD activity displayed a continuous increase in line with the weeks of lactation (which to our knowledge has not yet been studied) vitamin E might increase this activity, as seen from an *in vitro* study (28). The increase in lipid peroxidation measured thanks to plasma MDA levels, was consistent with other observations in dairy cows over the course of lactation up to 76 days (29). Our results are consistent with those of Bouwstra et al., (10, 30) who also observed that MDA levels fell in cows supplemented with vitamin E. Our assumption is that because vitamin E is close to the lipid membrane of cells, it increases cellular GPx activity compared to plasma GPx activity (31) and thus reduces lipid peroxidation (32). As for plant extracts, no effects were observed on the redox status of the cows except for a tendency toward lower MDA levels at week 12 of lactation; this is consistent with the previously described effects of another plant supplementation, i.e. green tea (33).

At the local scale of mammary tissue, we tended to observe an antioxidant effect with plant extracts but not with vitamin E. In week 4, the quantification of gene expression in milk MEC showed that the *NFE2L2* gene (involved in the Nrf2 pathway) tended to be more expressed in the plant extract group than in the vitamin E group. This is consistent with *in vitro* studies showing activation of the Nrf2 pathway with plant extracts (34–36). Thanks to their respective antioxidant functions, it is possible that vitamin E and plant extracts limit the increase in lipid peroxidation and oxidative stress, but the effects of vitamin E or plant extract supplements on cow redox status were scarce.

Considering immune status, the low-grade inflammation present between the start of lactation and peak lactation is associated with metabolic changes (2). In fact, at week 2 of lactation, a high plasma concentration of haptoglobin was observed, this being an acute inflammation protein. This was consistent with other observations (37), but without the supplements exerting any effects. We also observed a high white blood cell count at week 2 of lactation, consistent with other studies (38, 39), but again, no effect of the supplements. Indeed, during our study, the multiparous cows already had, or tended to have, higher monocyte, neutrophil and eosinophil counts than the primiparous cows in the control group. In multiparous cows, we also observed lower white blood cell, monocyte and eosinophil counts with the vitamin E and/or plant extract supplementations. One study had shown a decrease in granulocytes (including neutrophils, eosinophils, basophils) with plant compound supplementation at day 7 of lactation, but this study only extended until day 14 of lactation (40). We were able to show that antioxidant supplementation might offer a strategy to modulate cell counts according to the parity of the cows. Finally, vitamin E and plant extract supplementations increased the number of classical neutrophils and reduced that of regulatory neutrophils. It has been shown that regulatory neutrophils are able to suppress the T lymphocyte response *in vitro* (41, 42). Finally, vitamin E and plant extract supplementations may enable modulation of the immune response with a decrease of regulatory neutrophils, in such a way as to avoid immune suppression in early lactation.

Considering the systemic immune functional capacities of classical and regulatory neutrophils, vitamin E and plant extract supplementation reduced ROS production following *ex vivo* stimulation. These results were consistent with those of other studies at the scale of the whole neutrophil population (4, 43) but none of those studies differentiated the classical and regulatory populations. It is interesting to note that vitamin E and plant extract supplementation also affected plasma cytokine concentrations in the context of *ex vivo* stimulation with HK *E. coli*, causing a reduction in CCL4 (chemokine), and IL6 and IL8 (pro-inflammatory cytokines) levels. This was consistent with studies which had shown that vitamin E supplementation reduced inflammation and more specifically, levels of the pro-inflammatory cytokines IL8 (44, 45) and IL6 (42) in both humans and cows. It was also consistent with other studies showing that supplementation with tannin extracts or green tea lowered pro-inflammatory cytokine levels (IL8, IL1β and TNFα) (33, 46). It is interesting to highlight that in our study, the alleviating effects of the nutritional strategy on the inflammatory response could only be observed after *ex vivo* stimulation. Only two cytokines were more elevated in the vitamin E (IFNγ) and plant extract (IL17A and IFNγ) groups than in the control group. These two cytokines are produced by CD4^+^ T-lymphocytes to trigger leukocytosis during infection and permit an antigen-specific response in the mammary gland (47). As described during the same study, this immune response may limit the severity and sequelae of mastitis.

In milk MEC, our results suggest that vitamin E and plant extract supplementations affected immune response genes. Compared with the control group, at week 12, supplementation with vitamin E and plant extracts reduced *FABP3*, a protein known for its antimicrobial function (48). The transcriptomic analyses also revealed some specific effects of vitamin E versus plant extract with a downward trend in *CXCL10*, a decrease in *LBP* at week 8 in multiparous cows, and an increase in *IFN*β*1* at week 4 in primiparous cows. A specific effect was observed with plant extract supplementation in the form of a lower *LBP* level at week 12 compared with the control group in multiparous cows. These effects indicate the ability of nutritional supplementation to modify the immune response of MECs, particularly in multiparous cows. Further research is necessary to understand whether these dynamics of immune gene expression in the mammary gland – i.e. increasing at week 4 and then decreasing at week 12 – may suggest a better protection during the first months of lactation.

The nutritional supplementation did not modify SCC. In another study, vitamin E supplementation similar to ours was given to dairy cows, i.e. 3000 IU before and 1000 IU after calving and the milk SCC fell (11). A lower SCC has also been reported previously with a plant extract rich in antioxidant (49) and a citrus flavonoid extract (50). One reason for these discrepancies may be that the milk SCS was low in our experiment in all cows even those in the control group. It ranged from 28,000 cells/mL to around 200,000 cells/mL, which probably indicates good basal health status (51). In accordance with the absence of SCC changes, and as indicated by the milk Na^+^:K^+^ ratio and the exfoliation of MEC into milk, the supplements did not affect mammary epithelium integrity. However, modifications to the expression of a gene coding for a constitutive tight junction protein might indicate an improvement in mammary epithelium integrity. Indeed, primiparous cows exhibited a trend toward higher *TJP1* gene expression with vitamin E than with the control treatment, and we also observed that plant extract supplementation tended to increase *TJP1* in multiparous compared to primiparous cows. In a previous *in vitro* study, vitamin E had also been shown to limit damage to bovine mammary endothelial cells (52). No significant variations were observed in gene expression and content related to milk fat, protein synthesis in milk with vitamin E or plant extract supplementation. The only effect seen was a lower lactose content in the vitamin E and plant extract group of multiparous cows, without any specific difference in *LALBA* gene expression or mammary epithelium integrity. This effect could be linked to cow genetics, or to multiple infections they may have encountered during lactation (53).

## CONCLUSIONS

At the systemic level for vitamin E supplementation, and at the local mammary gland level for plant extract supplementation, we observed a tendency towards an improved antioxidant response, which may have resulted in reduced lipid peroxidation. After *ex vivo* stimulation, it appeared that immunity was differentially regulated, leading us to conclude that supplementation with vitamin E or plant extracts could prevent systemic hyper-inflammation. Our study also illustrates the need to take parity into account in order to better define an antioxidant supplementation strategy, because multiparous cows, supplemented with vitamin E or plant extract, appeared to revert to the physiology of primiparous cows. Vitamin E and plant extracts require further study so as to better understand their preventive action, which could be verified by studying their effects in inducing an inflammatory challenge.

## METHODS

### Animals, experimental design and animal housing

This experiment was conducted at the INRAE Experimental Farm (IE PL, INRAE, Dairy Nutrition and Physiology; https://doi.org/10.15454/yk9q-pf68, 35650 Le Rheu, Brittany, France accreditation for animal housing no. C-35–275-23”) between 23 August and 17 December 2021. The experiment was approved by the Rennes Ethics Committee on Animal Experimentation and the French Ministry for Higher Education, Research and Innovation (APAFIS project number #31835-2021053017243978_v) and performed in compliance with all applicable provisions established by European Directive 2010/63/EU. The study was built and reported in accordance with ARRIVE guidelines (https://arriveguidelines.org).

Fifty-one Holstein dairy cows were assigned to three groups of 17 cows with a final target of 15 cows per group, to overcome health disorder that could happen at calving. The groups were followed between 3 weeks before and 12 weeks after calving. All cows were fed with a total mixed ration (TMR) based on supplemented maize silage and formulated in accordance with INRAE guidelines (54). The TMR is described in Supplementary Table S6. Three supplementation strategies were assigned to each group. The ‘control’ group received only the standard TMR and included 15 cows, the ‘vitamin E’ group received the standard TMR supplemented with vitamin E and included 16 cows and the ‘plant extracts’ group received the standard TMR supplemented with plant extracts of Biodevas Laboratoires (Savigné-L’Evêque, France) and included 14 cows. The number of cows included in the study was based on a previous study involving 12 cows in order to evaluate the effect of vitamin E supplementation on immunity criteria (43). In order to limit the variability between groups, for multiparous cows, groups were balanced for the expected calving date, the lactation rank (control: 2.9 ± 0.7, vitamin E: 2.6 ± 0.7, plant extracts 3.0 ± 0.7), the milk production during the first 60 days of lactation of their previous lactation (control: 2020 ± 212 L, vitamin E: 2092 ± 193 L, plant extracts 2033 ± 209 L), their average milk somatic cell count (SCC) over the last month of the previous lactation (control: 5.7 ± 4 .10^3^ cell/mL, vitamin E: 7 ± 5 .10^3^ cell/mL, plant extracts 7.9 ± 4 cell/mL), and their last body condition score (BCS) (control: 2.1 ± 0.1, vitamin E: 2.0 ± 0.1, plant extracts 2.2 ± 0.1). For primiparous cows, groups were only balanced for the expected calving date. The cows were also grouped into 4 blocks according to calving date, with each block sharing a common calendar week of calving. To maintain a maximum interval of 7 days between calving dates of successive blocks, some cows were switched between blocks according to their actual calving date. We carefully balanced the parities within each group and block and we checked the blocks remained balanced with actual calving date. Six cows were excluded and reasons were: late calving, more than one week outside the trial after the 4^th^ blocks; hock problem; significant weight loss; lowered hip; abortion and accidental death. During the experiment, 2 benign mastitis cases were detected during blood sampling in control group (one on day 3 and one on week 2 of lactation), but these had no effect on milk production or blood data and did not induce outliers in the physiological data of interest. We also detected a metritis at week 2 of blood sampling for a cow in the vitamin E group, but none of the measured physiological parameters considered in this study were affected. These latter three cows were not excluded. The control group included eight multiparous and seven primiparous cows (N = 15), the vitamin E group included eight multiparous and eight primiparous cows (N = 16), and the plant extract group included eight multiparous and six primiparous cows (N= 14).

The cows were housed in a free stall barn with free access to feed and water with controlled individual ingestion, except during the 1 to 2 days around calving when they were kept in individual stalls. In the free stall barn, all cows had access to a unique individual trough and the barn was equipped with a robot weighing, for each cow, the offered amount of TMR and refusals. In the vitamin E group, all-rac-α-tocopheryl acetate (Cooperl, Plestan, France) was added to the diet in a solid form at a concentration of 30,000 mg/kg (1 mg/eq IU). Three weeks before calving, the cows were given 3000 IU/day (100 g/day), then between calving and week 12 of lactation they received 1000 IU/day (35 g/day) (11–13). The plant extracts given as a supplement to the corresponding group comprised *Sambucus nigra*, *Salix alba*, *Laurus nobilis, Haragophytum procumbens*, *Silybum marianum* and *Arctium lappa* (Biodevas Laboratoires, Savigné-L’Evêque, France). The plant extracts had undergone Afnor certification, which recognizes compliance of the product with ISO 22,000, i.e. safe and without risk to food safety. The plant extracts had also been certified under the Feed Certification scheme (GMP+ International) to ensure food safety worldwide. Plant extracts were given between calving and week 12 of lactation at a rate of 10 g/day in a solid form. The dosage of 10 g per day has been set according to a criteria of compatibility with industrial premix manufacturing processes. This dose is similar to that used for other plant extracts (49). Both vitamin E and plant extracts were top dressed on the TMR. The diet was distributed twice daily in 2 equal-sized meals, at 08:30 am and 5:00 pm. Feed distribution and feed refusals were measured in order to calculate DMI. The centesimal and chemical compositions of the diet are presented in detail in Supplementary Table 1. The cows were milked twice daily, at 07:00 am and 4:30 pm. Milk yield was recorded individually at each milking.

Health events were monitored daily and cows with health issues at calving (stillborn calf, abortion) or lameness problems requiring isolation from the herd, were excluded. The body condition score (BCS) of each cow was evaluated in weeks -2, 2, 8, and 12 approximately.

### Vitamin E and A, minerals and other forage analyses

Forage samples were analyzed for mineral concentrations (macro and trace elements) using ICP-OES (5110 Agilent Technology), as previously described (55). Levels in forage of γ- and α-tocopherols were quantified in duplicate using an Acquity UPLC system (Waters, Saint-Quentin-en-Yvelines, France), as previously described (56). The daily ingested amounts of vitamin E (α and γ) was measured by considering the DMI and their contents in the diet, with all-rac-α-tocopheryl acetate supplementation for the vitamin E supplemented group. The NE_L_ balance and DPI efficiency were calculated according to INRAE guidelines (54).

### Milk samples and determinations of milk composition

Milk samples were collected after morning milking as previously described for each of the 45 cows in week 4, 8 and 12 (19). Briefly, milk samples were analyzed weekly to determine their protein, fat and SCC content, in order to calculate the somatic cell score (SCS) (Mylab, Châteaugiron, France), with additional samples being collected in weeks 4, 8 and 12 from the entire morning milking to determine the lactose content (Mylab). At the same time points, a 1.8-liter milk sample was collected for the isolation of mammary epithelial cells (MEC) which are valid sources for mammary transcripts (57). Moreover, a 5 mL milk sample was also collected at the same time points to determine milk sodium and potassium concentrations as markers of mammary epithelium integrity (19).

### Blood samples

Blood samples were collected just after milk samples around 8:30 in the morning, as previously described (19). Briefly, they were collected from the coccygeal vein or jugular vein, depending on the type of analyzes to be performed, at week -3 (before supplementation), weeks -2 and 2 (for mineral and vitamin measures only), and weeks 4, 8 and 12. Samples from the coccygeal vein were placed in tubes containing lithium heparin as an anticoagulant (Vacutest, Kima srl, Arzergrande, Italy) and used to analyze the metabolic parameters, redox balance and inflammation markers detailed in the biochemical analysis section. Two other 9 mL tubes containing K_2_ EDTA as an anticoagulant (Vacutest) were used for the determination of MDA and vitamin E. Plasma was separated by centrifugation at 1,200 × *g* for 10 min at 4°C and stored at −20°C. Blood samples from the jugular vein were placed in two monovettes for plasma cytokines (S-Monovette, Sarstedt Marnay, France); one tube contained lithium heparin as an anticoagulant (Vacutest) to determine antioxidant enzymatic activity GPx, and another 9 mL tube contained K_2_ EDTA as an anticoagulant (Vacutest) to determine the blood neutrophil phenotype and to measure ROS production.

### Blood cytokine and chemokine measurements following *E. coli* stimulation

The immune response of cows was evaluated by measuring the plasma levels of a panel of 15 cytokines thanks to an *ex vivo* challenge using blood cells with or without heat-killed *E. coli* (HK *E. coli*). The blood samples in the two 1 mL-monovettes – one a control without bacteria and one containing HK *E. coli* (final concentration of 10^7^ *E. coli* P4 bacteria) – were incubated and treated as previously described (19, 58). The results were supplied in MFI (Median Florescence Intensity), and transformed into decimal logarithms.

### Hematological profile and flow cytometry

The hematological profile was analyzed by the Labocea laboratory (Fougères, France) at the same sample collection time points used to determine the white blood cell count (10^9^/L). The percentages of lymphocytes, monocytes, polynuclear neutrophils, and polynuclear eosinophils were multiplied by the total concentration of white blood cells, to obtain their concentration. Neutrophils and their ROS production stimulated or unstimulated by tert-butyl hydroperoxide (TBHP) were measured by flow cytometry as previously described, using a MACSQuant Analyzer 10 cytometer (Miltenyi Biotec) (19). The results were analyzed with Kaluza software 114 (analysis version 2.1, 2009-2021 Beckman Coulter, Inc.) with the results supplied in percent for neutrophil count (%) and their ROS production in MFI (Mean Fluorescence Intensity) (41).

### Mammary epithelial cell (MEC) count and gene expression

To determine the concentration and exfoliation rate of MEC in milk, and gene expression in milk MEC, milk cells were prepared from fresh milk samples collected at morning milking (1.8 kg in eight tubes of 233 g of milk), as previously described (19). Briefly, MEC were isolated from somatic cells using an immunomagnetic method. After defrosting at room temperature, milk MEC samples were crushed and homogenized in 1 mL TRIzol and ground mechanically with 2.8 mm metal beads (P000925-LYSK0-A.0, Ozyme, Saint-Cyr-l’École, France) using a Precellys ball mill (DQ2368, Bertin, Montigny-le-Bretonneux, France), followed by extraction using the RNeasy Mini kit (RNeasy Mini Kit QIAGEN, Hilden, Germany) (59). The amounts of total RNA extracted from milk MEC samples were determined using a DeNovix DS-11 Spectrophotometer (DeNovix Inc., Wilmington, DE). Total RNA quality was assessed with a Bioanalyzer with on average a RNA Integrity Number of 6.13 (Agilent Technologies, Massy, France) and 10-100 ng were reverse transcribed with the iScript Reverse Transcriptase mix (Bio-Rad, Hercules, CA94547, USA) according to the manufacturer’s instructions. Primers (Eurogentec, Seraing, Belgium) are listed in Supplementary Table S7. Primer validation was performed on a serially diluted pool of complementary DNA using a LightCycler 480 Real-Time PCR System (Roche, Boulogne-Billancourt, France). Firstly, pre-amplification was performed with steps of 2 min at 95°C, 20 cycles consisting of 15s - 95°C and 4 min -60°C to 4°C to infinity, for 1.25 µL complementary DNA and 3.75 µL of amplification solution. The pre-amplification solution for 96 genes comprised a mix containing all the forward and reverse primers (v/v, 13:100) (for 96 genes to be amplified, 1 µL of primer forward and 1 µL of primer reverse of 10 µM of stock solution). The volume of this mix was adjusted to 200 µL with RNase DNase free buffer (10 mM Tris, 8.0 pH, 0.1 mM EDTA, Teknova, CA, USA), a preAmp Master Mix solution (v/v, 27:100) (Fluidigm 100-5581) and ultra-pure water (v/v, 6:100). This pre-amplification was followed by exonuclease treatment for 30 min 37°C then 15 min 80°C, and then 4°C (exonuclease I, New England Biolabs). After the exonuclease reaction, the samples were diluted five-fold with a buffer (10 mM Tris-HCl, 1.0 mM EDTA, Teknova), and could then be stored at -20°C until Fluidigm quantification. This quantification was performed with Biomark HD (Fluidigm) in a 48×48 well plate, according to the manufacturer’s instructions. The annealing temperature was 60°C. Data were analyzed with Fluidigm Real-Time PCR software to determine the cycle threshold (Ct) values (41). The Ct values of milk MEC were then expressed relative to the geometric mean of four reference genes (*ACTB*, *GAPDH*, *PPIA* and *RPLP0*) using a semi-absolute method, as previously reported (60). Briefly, the approximate number of molecules of RNA (mol_nb_gene_) was calculated using the following formula mol_nb_gene_ = Ct_gene_ - 40 / -3.32, where 40 corresponding to the number of cycles and 3.32 is the slope corresponding to the maximum PCR efficiency of 100% for the primers; in parallel we calculated 10^geometric mean of Ct values of reference genes^ ; and finally the ratio of the two previous calculations to obtain the semi-absolute value for each gene of interest: log_10_((mol_nb_target_gene_ / mol_nb_reference_gene_) × 10^8^).

### Biochemical analysis: plasma metabolites, haptoglobin and redox status determination

Plasma glucose, urea, NEFA, BHB for energy metabolism, ROM for oxidative stress, BAP and FRAP for total antioxidant capacities, and haptoglobin for inflammation were measured using a multiparameter analyzer (Kone Instrument Corp., Espoo, Finland) and according to the kits as previously described (19). After antioxidant measurement, an oxidative stress index (OSI) was calculated with ROM on FRAP noted ISO_FRAP_, and with ROM on BAP noted ISO_BAP_ (23), and the activity of the GPx enzyme was measured in plasma and erythrocytes as previously described (19). Plasma calcium concentrations were obtained by ICP-OES (5110 Agilent Technology), as previously described (19). The plasma concentration of cortisol was assessed using the ELISA method. The ELISA plates were coated with 200 µL/well of mouse monoclonal antirabbit immunoglobulin antibody (Bertin Pharma, Montigny-le-Bretonneux, France), as previously described (61). For these manipulations, the CVs do not exceed 5% inter- and intra-assay.

Plasma concentrations of γ-tocopherol, α-tocopherol, β-carotenes and retinol were quantified in duplicate from plasma, as previously described (56), for this manipulation, the CVs do not exceed 25% inter- and intra-assay. Plasma MDA was determined by HPLC-fluorescence adapted from (62). Briefly, a solution of 50 μL butyl hydroxytoluene in ethanol (0.05%) and 50 μL plasma sample was added to 400 μL phosphoric acid (0.44 M in H_2_O) for deproteinization and vortexed, to which was added 100 μL 0.6 % thiobarbituric acid in H_2_O before incubation at 90°C for 1 h 30. After cooling in an ice bath and adding 500 μL *n*-butanol, the mixture was centrifuged at 10,000 × *g* for 5 min at room temperature to recover the MDA-TBA_2_ complex present in the upper phase. Twenty µL of this phase were injected into the HPLC System (Alliance, Waters, Guyancourt, France) equipped with an reverse phase C18 column (XBridge 3.5 μm, 4.6 × 150 mm; Waters, Guyancourt, France). Isocratic elution of the mobile phase contained dihydrogen phosphate buffer (0.05 M, pH 6.8) and methanol (60:40; v/v) with a flow rate set at 0.4 mL/min. The retention time (5.80 min) and the entire analysis time (7 min) of MDA-TBA_2_ was detected by fluorescence (excitation at 532 nm and emission at 553 nm), and its plasma concentration was determined using a 1,1,3,3-tetraethoxypropane (Merck, Darmstadt, Germany) calibration curve. For this manipulation, the CVs do not exceed 20% inter- and intra-assay.

### Statistical analysis

All data were analyzed using the lmerTest-package for a linear mixed effects model procedure on R Studio (version 1.3.1093, 2009-2020). According to the following statistical model, a type III ANOVA test was used:

Y (*ijklm*) = *µ* + covariate + parity *_i_* + treatment *_j_* + week *_k_* + (parity x week) i*_k_* + (treatment x week) *_jk_* + (parity x treatment) *_ij_* + (parity x treatment x week) *_ijk_* + cow *_l_* + block *_m_ +* □ *_ijklm._* Y _*ijklm*_ was the analyzed variable, parity *_i_* the fixed effect of parity, (i.e. primiparous or multiparous) (1 ddl), treatment *_j_* was the fixed effect of the group, i.e. control, vitamin E or plant extracts (2 ddl), week *_k_* was the fixed effect of sampling week relative to calving date (the number of ddl depending on the sampling numbers for each analyzed variable), cow *_l_* was the cows considered as a random effect, block *_m_* was a fixed effect of the blocks (3 ddl) and □ ijklm was the residual error. The data obtained from blood samples before supplementation (at week -3 relative to calving) were used as covariates for certain variables and were also included in the model, in order to highlight the effects of supplementation (vitamin E and plant extract groups). The same mixed model was used for the data obtained from the milk samples, but without a covariate value. The means adjusted from the model (emmeans) and SEM (standard error of the mean) were used to draw the graph. Post-hoc paired comparisons between modalities of parity, weeks, or interactions were also calculated according to Satterthwaite’s method. The thresholds of significance were set at ∗ *P* < 0.05; ∗∗ *P* < 0.01, and ∗∗∗ *P* < 0.001, and trends were noted at ^†^ *P* ≤0.10. The Figures were drawn using R Studio and were based on emmeans and SEM resulting from the double interactions and more specifically, the treatments according to the weeks relative to calving, or the treatments according to parity according to the Figures. Student’s t test was used only once when the multiparametric test failed to indicate an interaction between parity or week and treatment.

## Acknowledgements

The authors would like to thank Dr. Gaetan Vetea Plichart, all the technicians and animal staff at the IE PL Experimental Farm (le Rheu, France), an particularly, Gaël Boullet, Françoise Pichot and Severine Urvoix and the laboratory technicians in the PEGASE Joint Research Unit (Saint-Gilles, France): Perrine Poton, Maryline Lemarchand and Jacques Portanguen, for their assistance and support during the planning and execution of the experiment. Our thanks also go to the technicians, and in particular Christophe Gitton in the Bacterial Infections and Immunity of Ruminants (IBIR) team at the INRAE ISP unit (Nouzilly, France).

## Author contributions

Drs A.B., M.B., P.G., B.G. and A.R. designed the protocol and all the experiments in dairy cows, performed the experiments, participated in the statistical analysis and interpretation of the results, and wrote the manuscript. A.C. performed the experiments in dairy cows, performed certain laboratory analyses, created the database, performed statistical tests, interpreted and represented the results, and wrote the manuscript. Dr J-FR. provided funding, supported the design and implementation of the study, and reviewed the manuscript. O.D. assured the complete monitoring of the experimental farm and the collection of zootechnical data. C.M., S.P. and L.L-N designed a laboratory analysis by adapting the protocols to dairy cows. All authors read and approved the final manuscript.

## Additional information

Supplementary data are available at https://figshare.com/s/bcce1f9fd2542a7462fc?file=51895343. DOI: 10.6084/m9.figshare.27266931.

