## supplementary materials for "Nutritional supplementation with vitamin E or plant extracts affects redox and immune response in early lactating dairy cows"

**Table S1**: Plasma metabolic biomarkers in unsupplemented group (n = 15), vitamin E supplemented group (n = 16), and plant extract supplemented group (n = 14) dairy cows in day 1, and weeks 2, 4, 8 and 12 relative to calving. Metabolic biomarkers in plasma such as non-esterified fatty acids (NEFA), µmol/L, β-hydroxybutyrate (BHB), µmol/L, urea, mg/L, and glucose, mg/dL were measured from 2 to 12 weeks relative to calving, and day 1 for calcium concentration.

| Item [1] | Control | | | | | | Vitamin E | | | | | | Plant extracts | | | | | | *P*-value [2] | | |
|---|---|---|---|---|---|---|---|---|---|---|---|---|---|---|---|---|---|---|---|---|---|
| | Day 1 | 2 | 4 | 8 | 12 | SEM | Day 1 | 2 | 4 | 8 | 12 | SEM | Day 1 | 2 | 4 | 8 | 12 | SEM | TREAT | WK | TREAT x WK |
| NEFA (µmol/L) | NE[3] | 828 | 579 | 289 | 234 | 73.42 | NE | 812 | 488 | 299 | 214 | 71.13 | NE | 1047 | 738 | 389 | 294 | 83.51 | **≤ 0.10** | **< 0.001** | 0.8 |
| BHB (µmol/L) | NE | 787 [a] | 1043 | 761 | 795 | 138.8 | NE | 1334 [b] | 969 | 663 | 756 | 132.8 | NE | 840 [a] | 1355 | 649 | 605 | 153.5 | 0.74 | **< 0.001** | **< 0.05** |
| Urea (mg/L) | NE | 164 | 137 | 169 | 224 | 11.5 | NE | 170 | 130 | 171 | 222 | 10.93 | NE | 152 | 122 | 173 | 216 | 12.2 | 0.68 | **< 0.001** | 0.97 |
| Glucose (mg/dL) | NE | 68.4 | 66.6 | 72.8 | 73.8 | 1.672 | NE | 63.8 | 65.2 | 72.2 | 73.5 | 1.63 | NE | 68.8 | 65.2 | 72.7 | 76.2 | 1.786 | 0.49 | **< 0.001** | 0.27 |
| Calcium (mg/L) | 79.1 [a] | 88.9 | 87.4 | 89.1 | 89.8 | 2.505 | 81.5 | 88.3 | 92 | 90.8 | 94.9 | 2.412 | 88.1 [b] | 95.2 | 92.3 | 88.3 | 94.2 | 2.695 | 0.35 | **< 0.001** | **≤ 0.10** |

[1] Plasma samples were collected in day 1, and in weeks 2, 4, 8 and 12 relative to calving. The plasma sample in week -3 serves as a covariate because this plasma sample was collected before the supplementation with vitamin E or plant extracts.

[2] Probability: TREAT = effect of treatment (unsupplemented or vitamin E or plant extract group); WK = effect of day 1, week 2, 4, 8 and 12 relative to calving; TREAT × WK = the interaction between treatment and week. A covariate was chosen for the static calculations to highlight the effect of supplementation (vitamin E and plant extract group).

[3] NE =Not estimated because no collection.

**a-b** Mean values in the same row with different (P < 0.05) for the interaction between treatment and week relative to calving.

**Table S2**: Hematological profils in unsupplemented (n = 15), vitamin E supplemented (n = 16), and plant extracts supplemented (n = 14) dairy cows between week 2 and 12 relative to calving.

| Item [1] | Control | | | | | Vitamin E | | | | | Plant extracts | | | | | *P*-value [2] | | |
|---|---|---|---|---|---|---|---|---|---|---|---|---|---|---|---|---|---|---|
| | 2 | 4 | 8 | 12 | SEM | 2 | 4 | 8 | 12 | SEM | 2 | 4 | 8 | 12 | SEM | TREAT | WK | TREAT x WK |
| **Red blood cells (.$10^{12}$/L)** | 6.31 | 5.84 | 6.08 | 6.25 | 0.666 | 6.19 | 6 | 5.94 | 6.25 | 0.643 | 6.28 | 5.88 | 5.94 | 6.32 | 0.709 | 0.99 | **< 0.001** | 0.13 |
| **Platelets (.$10^9$/L)** | 461 | 499 | 478 | 510 | 40.2 | 429 | 443 | 418 | 480 | 38.7 | 435 | 451 | 514 | 504 | 42.3 | 0.42 | 0.3 | 0.88 |
| **Haemoglobin (.$10^9$/L)** | 101 | 90.3 | 91.4 | 91.8 | 1.86 | 96.6 | 92.1 | 88.5 | 91.2 | 1.81 | 99.33 | 92.9 | 89.75 | 93.62 | 1.96 | 0.61 | **< 0.001** | 0.47 |
| **Hematocrit (L/L)** | 0.29 | 0.26 | 0.27 | 0.27 | 0.006 | 0.28 | 0.27 | 0.26 | 0.27 | 0.005 | 0.3 | 0.27 | 0.27 | 0.28 | 0.006 | 0.62 | **< 0.001** | 0.32 |
| **Mean corpuscular volume (.$10^{-15}$/L)** | 46.5 | 45.6 | 45 | 44.1 | 0.316 | 46.7 | 46 | 44.8 | 44.2 | 0.307 | 47 | 45.4 | 44.3 | 43.6 | 0.34 | 0.71 | **< 0.001** | 0.1 |
| **Mean hemoglobin concentration (.$10^9$/L)** | 344[a] | 341 | 337 | 336 | 2.038 | 339 | 339 | 338 | 336 | 1.956 | 333[b] | 343 | 338 | 337 | 2.127 | 0.51 | **< 0.05** | **< 0.05** |
| **White blood cells (.$10^9$/L)** | 9.24 | 7.08 | 6.25 | 6.22 | 0.666 | 7.32 | 7.41 | 7.25 | 6.98 | 0.643 | 8.69 | 7.16 | 6.13 | 6.66 | 0.709 | 0.99 | **< 0.001** | 0.13 |
| **Lymphocytes (.$10^9$/L)** | 4.83 | 3.51 | 3.22 | 3.56 | 0.404 | 4.52 | 4.44 | 4.27 | 3.57 | 0.39 | 5.13 | 4.07 | 3.47 | 3.48 | 0.43 | 0.52 | **< 0.001** | 0.32 |
| **Monocytes (.$10^9$/L)** | 0.45 | 0.22 | 0.22 | 0.21 | 0.044 | 0.28 | 0.21 | 0.27 | 0.19 | 0.043 | 0.4 | 0.27 | 0.33 | 0.19 | 0.047 | 0.25 | **< 0.001** | 0.17 |
| **Neutrophils (.$10^9$/L)** | 3.42 | 2.68 | 2.23 | 2.13 | 0.403 | 2.13 | 2.21 | 2.21 | 2.77 | 0.389 | 2.91 | 2.42 | 1.88 | 2.75 | 0.426 | 0.74 | 0.1 | 0.16 |
| **Eosinophils (.$10^9$/L)** | 0.57 | 0.7 | 0.6 | 0.35 | 0.094 | 0.31 | 0.49 | 0.42 | 0.39 | 0.091 | 0.32 | 0.47 | 0.51 | 0.32 | 0.099 | 0.15 | **< 0.05** | 0.58 |
| **Basophils (.$10^7$/L)** | 0.36 | 0.09 | 0.93 | 0.68 | 0.526 | 1.52 | 0.36 | 0.85 | 1.13 | 0.506 | 1.08 | -0.01 | 1.77 | 0.75 | 0.552 | 0.45 | **< 0.10** | 0.76 |

[1] Plasma samples were collected in day 1, and in weeks 2, 4, 8 and 12 relative to calving. The plasma sample in week -3 serves as a covariate because this plasma sample was collected before the supplementation with vitamin E or plant extracts.

[2] Probability: TREAT = effect of treatment (unsupplemented or vitamin E or plant extract group); WK = week 2, 4, 8 and 12 relative to calving; TREAT × WK = the interaction between treatment and week. A covariate is chosen for the static calculations to highlight the effect of supplementation (vitamin E and plant extract group).

**a-b** Mean values in the same row with different (P < 0.05) for the interaction between treatment and week relative to calving.

**Table S3**: Measurement of 15 cytokines in plasma expressed in mean fluorescence intensities (MFI) without an ex vivo heat killed *Escherichia coli* (HK *E. coli*) challenge in unsupplemented (n = 15), vitamin E supplemented (n = 16), and plant extracts supplemented (n = 14) dairy cows in weeks 8 and 12 relative to calving. The rates of variation for each cytokines had been calculated to represent the difference between unsupplemented, vitamin E and plant extract groups according to the week of lactation.

| Item [1] | Weeks relative to calving | Control | | Vitamin E | | Plant extracts | | Overall mean | SEM [2] | *P* -value [3] | | |
|---|---|---|---|---|---|---|---|---|---|---|---|---|
| | | MFI mean | Rate of variation | MFI mean | Rate of variation | MFI mean | Rate of variation | | | TREAT | WK | TREAT x WK |
| | 8 | 1.765 | 0.024 | 1.711 | -0.007 | 1.785 | 0.036 | | | | | |
| total mean | 12 | 1.689 | -0.020 | 1.683 | -0.023 | 1.705 | -0.010 | 1.7229 | 0.0258 | 0.483 | 0.050 | 0.727 |
| **Chemokine cytokines** | | | | | | | | | | | | |
| CCL2 | 8 | 3.372 | 0.008 | 3.323 | -0.006 | 3.406 | 0.018 | | | | | |
| | 12 | 3.350 | 0.002 | 3.301 | -0.013 | 3.313 | -0.009 | 3.3444 | 0.0367 | 0.606 | 0.301 | 0.757 |
| CCL3 | 8 | 1.833 | 0.097 | 1.679 | 0.004 | 1.699 | 0.016 | | | | | |
| | 12 | 1.560 | -0.067 | 1.618 | -0.032 | 1.642 | -0.018 | 1.6718 | 0.0828 | 0.931 | 0.185 | 0.581 |
| CCL4 | 8 | 2.916 | 0.037 | 2.849 | 0.013 | 2.876 | 0.023 | | | | | |
| | 12 | 2.816 | 0.001 | 2.764 | -0.017 | 2.649 | -0.058 | 2.8117 | 0.0697 | 0.675 | 0.090 | 0.733 |
| CXCL10 | 8 | 2.186 | 0.001 | 2.158 | -0.012 | 2.247 | 0.029 | | | | | |
| | 12 | 2.162 | -0.010 | 2.190 | 0.003 | 2.163 | -0.010 | 2.1844 | 0.0335 | 0.862 | 0.374 | 0.252 |
| **Pro-inflammatory cytokines** | | | | | | | | | | | | |
| IFNg | 8 | 1.060 | -0.002 | 1.061 | 0.000 | 1.079 | 0.017 | | | | | |
| | 12 | 1.009 | -0.050 | 1.028 | -0.031 | 1.132 | 0.067 | 1.0617 | 0.0210 | 0.075 | 0.689 | 0.227 |
| IL17A | 8 | 0.939 | 0.007 | 0.930 | -0.003 | 0.937 | 0.004 | | | | | |
| | 12 | 0.923 | -0.011 | 0.929 | -0.005 | 0.940 | 0.008 | 0.9331 | 0.0148 | 0.923 | 0.784 | 0.891 |
| IL1a | 8 | 1.200 | -0.056 | 1.272 | 0.001 | 1.369 | 0.076 | | | | | |
| | 12 | 1.227 | -0.035 | 1.264 | -0.006 | 1.297 | 0.020 | 1.2713 | 0.0392 | 0.300 | 0.599 | 0.502 |
| IL1b | 8 | 0.863 | 0.012 | 0.835 | -0.021 | 0.864 | 0.014 | | | | | |
| | 12 | 0.833 | -0.023 | 0.857 | 0.005 | 0.864 | 0.013 | 0.8525 | 0.0214 | 0.837 | 0.925 | 0.701 |
| IL2 | 8 | 1.536 | 0.005 | 1.540 | 0.008 | 1.610 | 0.054 | | | | | |
| | 12 | 1.486 | -0.027 | 1.495 | -0.022 | 1.498 | -0.019 | 1.5274 | 0.0175 | 0.417 | **< 0.001** | 0.090 |
| IL6 | 8 | 1.538 | 0.036 | 1.493 | 0.005 | 1.521 | 0.024 | | | | | |
| | 12 | 1.445 | -0.027 | 1.450 | -0.023 | 1.463 | -0.015 | 1.4850 | 0.0130 | 0.596 | **< 0.001** | 0.228 |
| IL8 | 8 | 2.540 | 0.047 | 2.229 | -0.082 | 2.447 | 0.008 | | | | | |
| | 12 | 2.567 | 0.057 | 2.354 | -0.030 | 2.427 | 0.000 | 2.4273 | 0.0947 | 0.196 | 0.693 | 0.860 |
| TNFa | 8 | 1.448 | -0.006 | 1.472 | 0.011 | 1.578 | 0.083 | | | | | |
| | 12 | 1.371 | -0.059 | 1.381 | -0.052 | 1.490 | 0.023 | 1.4567 | 0.0410 | 0.241 | 0.021 | 0.986 |
| **Anti-inflammatory cytokines** | | | | | | | | | | | | |
| IL10 | 8 | 1.401 | -0.026 | 1.419 | -0.014 | 1.533 | 0.066 | | | | | |
| | 12 | 1.385 | -0.037 | 1.447 | 0.006 | 1.446 | 0.005 | 1.4384 | 0.0330 | 0.310 | 0.415 | 0.298 |
| IL1Ra | 8 | 2.354 | 0.097 | 2.202 | 0.025 | 2.260 | 0.053 | | | | | |
| | 12 | 2.027 | -0.056 | 2.044 | -0.048 | 1.994 | -0.071 | 2.1470 | 0.0495 | 0.606 | **< 0.001** | 0.493 |
| IL4 | 8 | 1.285 | 0.050 | 1.251 | 0.022 | 1.277 | 0.043 | | | | | |
| | 12 | 1.169 | -0.045 | 1.177 | -0.039 | 1.186 | -0.031 | 1.2242 | 0.0112 | 0.572 | **< 0.001** | 0.394 |

[1] The MFI mean and rates of variation are presented to observe the effect of treatment according without HK E. coli stimulation. The formula for the rate of variation was calculated as follows: Rate of variation = (MFI mean - overall mean) / overall mean.

[2] The overall means and standard errors of the mean (SEM) were calculated for each cytokine.

[3] Probability: TREAT = effect of treatment (unsupplemented or vitamin E or plant extract group); WK = week 8 and 12 relative to calving; TREAT × WK = the interaction between treatment and week. A covariate was chosen for the static calculations to highlight the effect of supplementation (vitamin E and plant extract group).

**Table S4**: Measurement of milk content and biomarkers of the mammary epithelium integrity in unsupplemented group (n = 15), vitamin E group (n = 16), and plant extracts group (n = 14) dairy cows between week 2 and 12 relative to calving. The milk content measured fat and protein milk in g/kg, fat and protein milk ratio, fat and protein content in g, and lactose milk in g/kg. The biomarkers of mammary epithelial integrity measured by milk somatic cell count (SCC), somatic cell score (SCS), mammary epithelial cell (MEC) concentration, MEC exfoliation, milk concentrations of $Na^+$, $K^+$, $Na^+$:$K^+$ ratio.

| Item [1] | Control | | | | | Vitamin E | | | | | Plant extracts | | | | | *P*-value | | | |
|---|---|---|---|---|---|---|---|---|---|---|---|---|---|---|---|---|---|---|---|
| | 2 | 4 | 8 | 12 | SEM | 2 | 4 | 8 | 12 | SEM | 2 | 4 | 8 | 12 | SEM | TREAT | PAR | WK | TREAT × WK |
| **Milk fat (g/kg)** | 43.3 | 40.4 | 39.5 | 39.8 | 1.231 | 46.2 | 42.5 | 40.4 | 41 | 1.297 | 49.6 | 44.1 | 40.5 | 41.5 | 1.525 | 0.24 | 0.07 | **< 0.001** | 0.25 |
| **Milk protein (g/kg)** | 34.1 | 28.8 | 28.2 | 29.8 | 0.489 | 34.7 | 28.2 | 28 | 29.9 | 0.505 | 32.6 | 28.2 | 28.4 | 29.7 | 0.588 | 0.93 | 0.8 | **< 0.001** | 0.15 |
| **Milk fat:protein** | 1.28 | 1.41 | 1.41 | 1.34 | 0.042 | 1.34 | 1.51 | 1.45 | 1.37 | 0.044 | 1.52 | 1.58 | 1.43 | 1.41 | 0.041 | 0.13 | **< 0.05** | **< 0.001** | 0.14 |
| **Fat yield (g/d)** | 557 | 576 | 597 | 646 | 26.68 | 582 | 590 | 619 | 657 | 27.19 | 658 | 613 | 626 | 679 | 31.44 | 0.57 | **< 0.001** | **< 0.001** | 0.52 |
| **Protein yield (g/d)** | 456 | 419 | 434 | 491 | 16.64 | 447 | 405 | 436 | 490 | 16.74 | 457 | 407 | 447 | 499 | 19.22 | 0.9 | **< 0.001** | **<0.001** | 0.99 |
| **Milk lactose (g/kg)** | NE[5] | 50.5 | 50.5 | 51.1 | 0.370 | NE | 48.9 | 49.4 | 49.4 | 0.37 | NE | 49.1 | 49.3 | 50.1 | 0.420 | **< 0.01** | **< 0.001** | **< 0.001** | 0.55 |
| **Lactose yield (g/kg)** | NE | 736 | 779 | 857 | 28.12 | NE | 705 | 773 | 802 | 28.22 | NE | 706 | 765 | 853 | 32.00 | 0.65 | **< 0.001** | **< 0.001** | 0.48 |
| **SCC ($.10^3$ /mL)** | 235 | 68.8 | 70.6 | 28.8 | 75.73 | 327 | 159 | 74.1 | 40 | 81.23 | 169 | 151 | 70.4 | 103 | 96.53 | 0.7 | 0.5 | **< 0.001** | 0.97 |
| **SCS ($log_2$(SCC/100)+3)** | 1.56 | 0.67 | 0.79 | 1.56 | 0.440 | 1.4 | 0.85 | 0.8 | 1.4 | 0.44 | 2.1 | 1.27 | 1.29 | 2.1 | 0.500 | 0.68 | 0.53 | **< 0.001** | 0.99 |
| **MEC count [3] ($.10^3$ /mL)** | NE | 6.45 | 2.62 | 2.78 | 1.556 | NE | 6.83 | 4.2 | 2.54 | 1.494 | NE | 11.4 | 4.47 | 4.46 | 1.645 | 0.27 | 0.43 | **< 0.001** | 0.37 |
| **MEC exfoliation[4] ($.10^6$)** | NE | 123 | 52 | 59 | 34.03 | NE | 122 | 91 | 46 | 32.66 | NE | 200 | 93 | 84 | 36.04 | 0.49 | 0.09 | **< 0.001** | 0.44 |
| **Milk $Na^+$ (mg/kg)** | NE | 322 | 302 | 299 | 14.15 | NE | 336 | 319 | 321 | 13.56 | NE | 354 | 324 | 312 | 15.03 | 0.16 | 0.51 | **< 0.001** | 0.39 |
| **Milk $K^+$ ($.10^1$ mg/kg)** | NE | 177 | 178 | 180 | 4.160 | NE | 177 | 177 | 180 | 4.084 | NE | 171 | 172 | 172 | 4.525 | 0.47 | 0.38 | 0.49 | 0.98 |
| **Milk $Na^+$:$K^+$ ratio ($.10^{-3}$)** | NE | 181 | 169 | 166 | 8.007 | NE | 190 | 181 | 180 | 7.676 | NE | 209 | 190 | 182 | 8.504 | 0.16 | 0.72 | **< 0.001** | 0.39 |
| **Plasma lactose (mg/L)** | NE | 49.9 | 45.7 | 41.9 | 4.410 | NE | 47 | 36.2 | 41.4 | 4.237 | NE | 45.6 | 47.1 | 49.8 | 4.662 | 0.44 | 0.27 | 0.26 | 0.23 |

[1] Milking milk samples were collected for measurement of somatic and epithelial cell levels in week 4, 8, and 12 relative to calving. Milk minerals and plasma lactose were measured at the same week of lactation.

[2] Probability of parity effects: TREAT = effects of treatment, unsupplemented vs. vitamin E vs. plant extracts; PAR = effect of parity, primiparous vs. multiparous; WK = effect of week in week 2, 4, 8 and 12 relative to calving; TREAT × WK = interaction between treatment and week.

[3] MEC count was calculated with MEC count on milk sample volume (milk 1.8 kg) ratio.

[4] MEC exfoliation was calculated with MEC count (mammary epithelial cells/mL) on milk yield (kg) ratio.

[5] NE =Not estimated because no collection.

**Table S5**: Abundance of antioxidant function mRNA, immune function mRNA, and milk production mRNA determined by real-time quantitative RT-PCR in milk isolated mammary epithelial cells (MEC) in unsupplemented (n = 15), vitamin E supplemented (n = 16), and plant extracts supplemented (n = 14) dairy cows according to weeks 4, 8 and 12 relative to calving or parity, primiparous or multiparous cows.

| Item[1] | Parity | Control | | | | Vitamin E | | | | Plant extracts | | | | *P*-value[2] | | | | | | |
|---|---|---|---|---|---|---|---|---|---|---|---|---|---|---|---|---|---|---|---|---|
| | | 4 | 8 | 12 | SEM | 4 | 8 | 12 | SEM | 4 | 8 | 12 | SEM | PAR | TREAT | WK | TREAT × PAR | PAR × WK | TREAT × WK | PAR × TREAT × WK |
| ABCG2 | multiparous | 6.23 | 6.29 | 6.73 | 0.191 | 6.32 | 6.06 | 6.08 | 0.189 | 6.48 | 6.36 | 6.31 | 0.195 | | | | | | | |
| | primiparous | 6.84 | 6.55 | 6.97 | 0.216 | 7.26 | 6.45 | 6.86 | 0.200 | 6.68 | 6.57 | 7.17 | 0.223 | 0.001 | 0.775 | 0.003 | 0.495 | 0.139 | 0.060 | 0.194 |
| AHR | multiparous | 5.91 | 5.96 | 6.12 | 0.136 | 5.88 | 5.98 | 6.07 | 0.135 | 6.02 | 6.15 | 6.13 | 0.139 | | | | | | | |
| | primiparous | 5.88 | 5.87 | 6.24 | 0.154 | 5.93 | 5.81 | 6.06 | 0.143 | 5.95 | 5.91 | 6.24 | 0.158 | 0.668 | 0.596 | 0.002 | 0.953 | 0.182 | 0.917 | 0.928 |
| AOAH | multiparous | 6.24 | 6.28 | 6.49 | 0.198 | 6.11 | 6.09 | 6.35 | 0.196 | 6.13 | 6.26 | 6.31 | 0.203 | | | | | | | |
| | primiparous | 5.92 | 6.08 | 6.30 | 0.221 | 6.03 | 5.73 | 6.02 | 0.206 | 6.08 | 5.92 | 6.29 | 0.232 | 0.139 | 0.609 | 0.007 | 0.943 | 0.640 | 0.740 | 0.712 |
| BAX | multiparous | 6.28 | 6.27 | 6.27 | 0.147 | 6.13 | 6.23 | 6.22 | 0.145 | 6.36 | 6.43 | 6.24 | 0.150 | | | | | | | |
| | primiparous | 6.13 | 5.96 | 6.12 | 0.165 | 5.89 | 5.99 | 6.11 | 0.154 | 6.19 | 5.96 | 6.31 | 0.172 | 0.051 | 0.452 | 0.549 | 0.999 | 0.124 | 0.745 | 0.690 |
| CASP1 | multiparous | 6.31 | 6.14 | 6.45 | 0.151 | 6.05 | 6.23 | 6.40 | 0.150 | 6.36 | 6.48 | 6.32 | 0.155 | | | | | | | |
| | primiparous | 6.16 | 6.00 | 6.23 | 0.169 | 6.15 | 5.75 | 6.33 | 0.157 | 6.18 | 5.89 | 6.36 | 0.177 | 0.082 | 0.678 | 0.001 | 0.936 | 0.015 | 0.600 | 0.121 |
| CASP13 | multiparous | 7.42 | 7.22 | 7.53 | 0.182 | 7.36 | 7.33 | 7.48 | 0.180 | 7.51 | 7.48 | 7.34 | 0.185 | | | | | | | |
| | primiparous | 7.26 | 6.89 | 7.33 | 0.207 | 7.27 | 6.92 | 6.57 | 0.192 | 7.23 | 6.89 | 7.29 | 0.212 | 0.004 | 0.532 | 0.076 | 0.645 | 0.352 | 0.304 | 0.140 |
| CASP8 | multiparous | 6.39 | 6.32 | 6.62 | 0.128 | 6.26 | 6.26 | 6.49 | 0.127 | 6.48 | 6.50 | 6.30 | 0.131 | | | | | | | |
| | primiparous | 6.30 | 6.22 | 6.47 | 0.144 | 6.50 | 6.19 | 6.42 | 0.134 | 6.33 | 6.24 | 6.71 | 0.149 | 0.757 | 0.785 | 0.002 | 0.755 | 0.192 | 0.804 | 0.018 |
| CAT | multiparous | 6.82 | 6.78 | 7.10 | 0.108 | 6.72 | 6.88 | 7.03 | 0.107 | 6.92 | 7.02 | 6.93 | 0.110 | | | | | | | |
| | primiparous | 6.73 | 6.72 | 7.04 | 0.123 | 6.90 | 6.68 | 6.98 | 0.114 | 6.81 | 6.67 | 7.15 | 0.126 | 0.379 | 0.779 | 0.001 | 0.932 | 0.079 | 0.894 | 0.148 |
| CCL2 | multiparous | 8.12 | 8.07 | 8.23 | 0.077 | 8.20 | 8.17 | 8.17 | 0.077 | 8.17 | 8.08 | 8.14 | 0.078 | | | | | | | |
| | primiparous | 8.07 | 7.92 | 8.04 | 0.089 | 8.17 | 7.89 | 8.13 | 0.082 | 8.13 | 7.92 | 8.01 | 0.090 | 0.005 | 0.538 | 0.008 | 0.968 | 0.249 | 0.889 | 0.772 |
| CCL20 | multiparous | 6.96 | 6.79 | 7.18 | 0.186 | 6.99 | 6.93 | 7.05 | 0.184 | 7.29 | 7.11 | 6.96 | 0.190 | | | | | | | |
| | primiparous | 6.88 | 6.36 | 6.80 | 0.208 | 6.90 | 6.25 | 6.74 | 0.194 | 6.85 | 6.30 | 6.95 | 0.217 | 0.008 | 0.809 | 0.001 | 0.937 | 0.013 | 0.892 | 0.243 |
| CCL5 | multiparous | 6.37 | 6.27 | 6.60 | 0.175 | 6.32 | 6.21 | 6.38 | 0.174 | 6.41 | 6.35 | 6.35 | 0.179 | | | | | | | |
| | primiparous | 6.15 | 6.14 | 6.29 | 0.198 | 6.06 | 5.88 | 6.19 | 0.184 | 6.25 | 6.15 | 6.58 | 0.205 | 0.126 | 0.429 | 0.026 | 0.741 | 0.687 | 0.992 | 0.667 |
| CD14 | multiparous | 7.51 | 7.55 | 7.78 | 0.133 | 7.29 | 7.53 | 7.68 | 0.132 | 7.36 | 7.63 | 7.33 | 0.136 | | | | | | | |
| | primiparous | 7.09 | 7.19 | 7.53 | 0.151 | 6.96 | 7.14 | 7.50 | 0.140 | 7.41 | 7.15 | 7.62 | 0.155 | 0.009 | 0.637 | 0.001 | 0.363 | 0.025 | 0.208 | 0.256 |
| CD36 | multiparous | 7.87 | 7.90 | 7.72 | 0.073 | 7.89 | 7.95 | 7.71 | 0.072 | 7.87 | 7.84 | 7.72 | 0.074 | | | | | | | |
| | primiparous | 7.83 | 7.80 | 7.64 | 0.083 | 8.03 | 7.82 | 7.66 | 0.077 | 7.89 | 7.81 | 7.73 | 0.085 | 0.526 | 0.607 | < 0.001 | 0.775 | 0.285 | 0.622 | 0.705 |
| CTSB | multiparous | 7.67 | 7.75 | 1.94 | 0.455 | 7.43 | 7.64 | 2.65 | 0.452 | 6.53 | 7.53 | 1.64 | 0.461 | | | | | | | |
| | primiparous | 7.21 | 7.34 | 0.92 | 0.523 | 7.07 | 7.38 | 0.97 | 0.483 | 7.47 | 7.43 | 1.03 | 0.528 | 0.064 | 0.665 | < 0.001 | 0.342 | 0.098 | 0.891 | 0.841 |
| CDH1 | multiparous | 5.86 | 5.99 | 6.31 | 0.145 | 5.78 | 5.99 | 6.03 | 0.143 | 5.96 | 6.14 | 6.17 | 0.148 | | | | | | | |
| | primiparous | 5.75 | 5.84 | 6.40 | 0.163 | 6.12 | 6.08 | 6.49 | 0.152 | 5.58 | 5.66 | 6.44 | 0.169 | 0.873 | 0.742 | < 0.001 | 0.116 | 0.003 | 0.492 | 0.399 |
| CLDN1 | multiparous | 5.90 | 6.07 | 6.23 | 0.178 | 5.98 | 6.16 | 6.08 | 0.176 | 6.35 | 6.35 | 6.32 | 0.182 | | | | | | | |
| | primiparous | 5.67 | 5.66 | 6.38 | 0.200 | 5.85 | 6.08 | 6.43 | 0.186 | 5.81 | 5.34 | 6.23 | 0.208 | 0.069 | 0.726 | 0.001 | 0.163 | 0.002 | 0.141 | 0.673 |
| CLEC4A | multiparous | 7.18 | 7.14 | 7.32 | 0.161 | 6.95 | 7.16 | 7.26 | 0.160 | 7.10 | 7.31 | 7.21 | 0.166 | | | | | | | |
| | primiparous | 6.96 | 6.79 | 7.17 | 0.179 | 7.02 | 6.93 | 7.09 | 0.167 | 7.10 | 6.68 | 7.06 | 0.189 | 0.089 | 0.987 | 0.012 | 0.837 | 0.019 | 0.617 | 0.462 |
| CLEC7A | multiparous | 6.60 | 6.34 | 6.93 | 0.187 | 6.20 | 6.53 | 6.75 | 0.185 | 6.58 | 6.71 | 6.61 | 0.192 | | | | | | | |
| | primiparous | 6.20 | 5.88 | 6.25 | 0.207 | 6.38 | 6.12 | 6.46 | 0.193 | 6.48 | 6.11 | 6.63 | 0.219 | 0.031 | 0.663 | 0.001 | 0.562 | 0.027 | 0.253 | 0.109 |
| CSN1S1 | multiparous | 4.55 | 4.81 | 8.16 | 0.435 | 4.38 | 3.80 | 7.01 | 0.432 | 5.49 | 4.45 | 8.05 | 0.441 | | | | | | | |
| | primiparous | 5.28 | 5.04 | 8.47 | 0.500 | 5.41 | 4.64 | 8.67 | 0.462 | 5.20 | 4.90 | 8.30 | 0.506 | 0.016 | 0.251 | < 0.001 | 0.185 | 0.857 | 0.776 | 0.775 |
| CSN3 | multiparous | 8.29 | 8.39 | 8.20 | 0.196 | 8.55 | 8.22 | 8.17 | 0.195 | 8.51 | 8.22 | 8.37 | 0.199 | | | | | | | |
| | primiparous | 8.28 | 8.66 | 8.43 | 0.224 | 8.10 | 8.57 | 8.05 | 0.207 | 8.01 | 8.44 | 8.03 | 0.228 | 0.718 | 0.681 | 0.146 | 0.467 | 0.022 | 0.873 | 0.806 |
| CXCL10 | multiparous | 5.88 | 5.47 | 5.79 | 0.193 | 5.92 | 5.54 | 5.57 | 0.192 | 6.16 | 5.86 | 5.43 | 0.196 | | | | | | | |
| | primiparous | 6.26 | 6.03 | 6.08 | 0.220 | 5.77 | 5.35 | 5.74 | 0.204 | 5.85 | 6.08 | 6.31 | 0.225 | 0.082 | 0.067 | 0.058 | 0.230 | 0.077 | 0.591 | 0.129 |
| CXCL2 | multiparous | 8.00 | 7.92 | 7.95 | 0.104 | 7.96 | 7.88 | 7.99 | 0.104 | 8.02 | 8.05 | 7.98 | 0.106 | | | | | | | |
| | primiparous | 7.86 | 7.52 | 7.88 | 0.119 | 7.66 | 7.67 | 7.74 | 0.110 | 7.96 | 7.50 | 7.89 | 0.122 | 0.002 | 0.547 | 0.008 | 0.952 | 0.047 | 0.666 | 0.220 |
| CXCL8 | multiparous | 8.02 | 8.10 | 7.95 | 0.140 | 8.28 | 8.14 | 8.24 | 0.140 | 8.17 | 8.13 | 8.30 | 0.142 | | | | | | | |
| | primiparous | 7.99 | 7.83 | 8.17 | 0.162 | 7.85 | 8.04 | 7.59 | 0.149 | 8.02 | 7.89 | 7.88 | 0.163 | 0.001 | 0.830 | 0.895 | 0.095 | 0.856 | 0.674 | 0.126 |
| DEFB5 | multiparous | 6.31 | 6.28 | 6.39 | 0.314 | 7.01 | 7.08 | 7.19 | 0.310 | 7.01 | 6.99 | 6.82 | 0.324 | | | | | | | |
| | primiparous | 7.06 | 6.88 | 7.10 | 0.339 | 6.78 | 6.41 | 6.80 | 0.316 | 6.83 | 6.51 | 6.93 | 0.369 | 0.921 | 0.767 | 0.042 | 0.182 | 0.055 | 0.922 | 0.595 |
| DHX58 | multiparous | 6.48 | 6.06 | 6.58 | 0.187 | 6.09 | 5.95 | 6.31 | 0.185 | 6.29 | 6.19 | 6.27 | 0.192 | | | | | | | |
| | primiparous | 6.03 | 5.66 | 6.09 | 0.208 | 6.17 | 5.61 | 6.22 | 0.194 | 6.06 | 5.58 | 6.35 | 0.218 | 0.044 | 0.835 | 0.000 | 0.584 | 0.128 | 0.988 | 0.288 |
| FABP3 | multiparous | 7.56 | 7.58 | 7.59 | 0.128 | 7.58 | 7.42 | 7.13 | 0.127 | 7.79 | 7.63 | 7.12 | 0.131 | | | | | | | |
| | primiparous | 8.03 | 7.83 | 7.60 | 0.144 | 8.10 | 7.69 | 7.63 | 0.134 | 7.85 | 7.86 | 7.72 | 0.149 | 0.002 | 0.547 | < 0.001 | 0.656 | 0.558 | 0.228 | 0.017 |
| FAS | multiparous | 6.34 | 6.23 | 6.76 | 0.187 | 6.19 | 6.23 | 6.42 | 0.185 | 6.30 | 6.34 | 6.44 | 0.191 | | | | | | | |
| | primiparous | 5.89 | 5.70 | 6.22 | 0.209 | 5.96 | 5.93 | 6.20 | 0.194 | 5.88 | 5.76 | 6.31 | 0.218 | 0.006 | 0.972 | 0.001 | 0.713 | 0.522 | 0.633 | 0.786 |
| GPX1 | multiparous | 7.32 | 7.33 | 7.75 | 0.112 | 7.21 | 7.38 | 7.60 | 0.111 | 7.35 | 7.49 | 7.52 | 0.115 | | | | | | | |
| | primiparous | 7.22 | 7.23 | 7.56 | 0.125 | 7.39 | 7.25 | 7.59 | 0.117 | 7.33 | 7.17 | 7.72 | 0.131 | 0.474 | 0.945 | < 0.001 | 0.752 | 0.069 | 0.881 | 0.110 |
| GPX3 | multiparous | 6.29 | 6.03 | 6.25 | 0.197 | 5.89 | 6.02 | 5.99 | 0.195 | 6.20 | 6.14 | 6.11 | 0.201 | | | | | | | |
| | primiparous | 5.94 | 5.46 | 5.56 | 0.222 | 6.03 | 5.57 | 5.77 | 0.206 | 5.57 | 5.69 | 5.70 | 0.230 | 0.004 | 0.964 | 0.179 | 0.469 | 0.490 | 0.530 | 0.431 |
| HSPA8 | multiparous | 7.95 | 8.01 | 8.16 | 0.065 | 7.89 | 7.95 | 8.08 | 0.064 | 7.94 | 8.05 | 8.11 | 0.066 | | | | | | | |
| | primiparous | 7.88 | 7.92 | 8.23 | 0.074 | 7.95 | 8.03 | 8.13 | 0.068 | 7.81 | 7.87 | 8.07 | 0.075 | 0.480 | 0.609 | < 0.001 | 0.217 | 0.369 | 0.548 | 0.738 |
| IFNAR1 | multiparous | 6.85 | 6.77 | 7.19 | 0.145 | 6.65 | 6.73 | 6.97 | 0.143 | 6.76 | 6.82 | 6.86 | 0.148 | | | | | | | |
| | primiparous | 6.52 | 6.44 | 6.69 | 0.161 | 6.58 | 6.53 | 6.70 | 0.150 | 6.68 | 6.42 | 6.75 | 0.169 | 0.017 | 0.911 | 0.001 | 0.688 | 0.391 | 0.669 | 0.565 |
| IFNAR2 | multiparous | 6.60 | 6.55 | 7.00 | 0.162 | 6.51 | 6.50 | 6.69 | 0.160 | 6.62 | 6.62 | 6.66 | 0.166 | | | | | | | |
| | primiparous | 6.32 | 6.26 | 6.53 | 0.180 | 6.32 | 6.19 | 6.48 | 0.167 | 6.46 | 6.14 | 6.65 | 0.189 | 0.028 | 0.769 | 0.001 | 0.897 | 0.438 | 0.763 | 0.394 |
| IFNB1 | multiparous | 7.11 | 6.62 | 6.96 | 0.185 | 6.72 | 6.58 | 6.74 | 0.184 | 6.98 | 6.79 | 6.49 | 0.190 | | | | | | | |
| | primiparous | 6.57 | 6.36 | 6.55 | 0.209 | 6.70 | 6.21 | 6.83 | 0.194 | 6.06 | 6.17 | 6.70 | 0.217 | 0.014 | 0.601 | 0.005 | 0.471 | 0.019 | 0.499 | 0.030 |
| IL1A | multiparous | 7.85 | 7.77 | 5.81 | 0.199 | 7.69 | 7.73 | 6.15 | 0.197 | 7.52 | 7.69 | 6.20 | 0.201 | | | | | | | |
| | primiparous | 7.48 | 7.21 | 5.54 | 0.228 | 7.42 | 7.28 | 5.78 | 0.211 | 7.67 | 7.38 | 5.24 | 0.231 | 0.001 | 0.842 | < 0.001 | 0.988 | 0.284 | 0.669 | 0.284 |
| IL1B | multiparous | 8.08 | 8.03 | 5.99 | 0.197 | 7.97 | 7.92 | 6.32 | 0.196 | 7.72 | 7.90 | 6.36 | 0.199 | | | | | | | |
| | primiparous | 7.80 | 7.54 | 5.69 | 0.227 | 7.84 | 7.44 | 5.93 | 0.209 | 7.98 | 7.60 | 5.39 | 0.229 | 0.002 | 0.804 | < 0.001 | 0.994 | 0.093 | 0.661 | 0.283 |
| IL10 | multiparous | 6.62 | 6.43 | 6.86 | 0.186 | 6.56 | 6.49 | 6.69 | 0.184 | 6.70 | 6.76 | 6.57 | 0.191 | | | | | | | |
| | primiparous | 6.42 | 6.05 | 6.43 | 0.207 | 6.45 | 6.05 | 6.35 | 0.193 | 6.55 | 6.05 | 6.61 | 0.218 | 0.029 | 0.802 | 0.001 | 0.983 | 0.039 | 0.788 | 0.217 |
| IL2RA | multiparous | 5.63 | 5.60 | 6.04 | 0.235 | 5.43 | 5.38 | 5.64 | 0.233 | 5.60 | 5.69 | 5.91 | 0.242 | | | | | | | |
| | primiparous | 5.25 | 5.12 | 5.61 | 0.261 | 5.49 | 5.05 | 5.65 | 0.244 | 5.48 | 5.09 | 5.65 | 0.275 | 0.104 | 0.793 | 0.001 | 0.688 | 0.180 | 0.920 | 0.928 |
| IL33 | multiparous | 4.97 | 4.78 | 5.28 | 0.189 | 5.13 | 4.71 | 5.00 | 0.188 | 5.51 | 5.14 | 4.99 | 0.192 | | | | | | | |
| | primiparous | 5.26 | 4.90 | 5.27 | 0.216 | 5.79 | 4.80 | 5.16 | 0.200 | 5.30 | 4.70 | 5.37 | 0.220 | 0.314 | 0.802 | 0.001 | 0.404 | 0.235 | 0.226 | 0.155 |
| IL6 | multiparous | 6.62 | 6.47 | 6.61 | 0.191 | 6.68 | 6.45 | 6.68 | 0.189 | 6.95 | 6.72 | 6.79 | 0.195 | | | | | | | |
| | primiparous | 6.42 | 5.96 | 6.23 | 0.217 | 6.34 | 5.67 | 6.22 | 0.201 | 6.50 | 5.87 | 6.48 | 0.223 | 0.001 | 0.345 | 0.001 | 0.813 | 0.099 | 0.933 | 0.924 |
| IRF3 | multiparous | 6.22 | 6.16 | 6.56 | 0.111 | 6.10 | 6.22 | 6.42 | 0.110 | 6.06 | 6.30 | 6.38 | 0.113 | | | | | | | |
| | primiparous | 5.95 | 6.11 | 6.41 | 0.126 | 6.13 | 6.05 | 6.37 | 0.117 | 6.13 | 6.09 | 6.64 | 0.129 | 0.396 | 0.822 | < 0.001 | 0.556 | 0.335 | 0.861 | 0.220 |
| ITGA4 | multiparous | 6.31 | 6.26 | 6.55 | 0.224 | 6.22 | 5.78 | 6.43 | 0.222 | 6.30 | 6.52 | 6.19 | 0.228 | | | | | | | |
| | primiparous | 6.22 | 6.10 | 6.43 | 0.253 | 6.38 | 6.17 | 6.41 | 0.235 | 6.43 | 6.08 | 6.60 | 0.261 | 0.844 | 0.773 | 0.036 | 0.694 | 0.714 | 0.668 | 0.223 |
| LALBA | multiparous | 8.08 | 8.11 | 7.70 | 0.800 | 8.48 | 8.09 | 7.70 | 0.794 | 8.24 | 7.13 | 8.20 | 0.813 | | | | | | | |
| | primiparous | 6.03 | 7.67 | 7.15 | 0.916 | 3.21 | 7.71 | 7.37 | 0.847 | 5.16 | 6.85 | 4.96 | 0.931 | 0.001 | 0.483 | 0.062 | 0.551 | 0.003 | 0.435 | 0.151 |
| LAP3 | multiparous | 7.17 | 7.04 | 7.08 | 0.275 | 7.03 | 7.03 | 7.42 | 0.273 | 6.76 | 7.19 | 7.17 | 0.279 | | | | | | | |
| | primiparous | 7.02 | 6.48 | 6.58 | 0.314 | 6.91 | 6.55 | 6.90 | 0.291 | 7.15 | 6.79 | 6.27 | 0.320 | 0.027 | 0.886 | 0.575 | 0.975 | 0.069 | 0.418 | 0.738 |
| LBP | multiparous | 4.35 | 4.54 | 4.86 | 0.243 | 4.48 | 4.05 | 4.17 | 0.241 | 5.04 | 4.79 | 4.04 | 0.246 | | | | | | | |
| | primiparous | 5.36 | 4.60 | 5.08 | 0.279 | 5.45 | 4.82 | 5.32 | 0.258 | 5.09 | 4.58 | 5.54 | 0.282 | 0.001 | 0.689 | 0.021 | 0.153 | 0.033 | 0.779 | 0.030 |
| LCN2 | multiparous | 5.77 | 5.71 | 5.97 | 0.195 | 5.90 | 5.36 | 5.42 | 0.193 | 6.24 | 5.65 | 5.53 | 0.199 | | | | | | | |
| | primiparous | 6.02 | 5.67 | 5.98 | 0.220 | 6.40 | 5.69 | 5.85 | 0.205 | 5.83 | 5.28 | 5.84 | 0.228 | 0.379 | 0.734 | 0.001 | 0.210 | 0.327 | 0.097 | 0.286 |
| LPL | multiparous | 6.53 | 6.59 | 6.63 | 0.134 | 6.59 | 6.70 | 6.46 | 0.133 | 6.74 | 6.86 | 6.37 | 0.137 | | | | | | | |
| | primiparous | 6.69 | 6.58 | 6.53 | 0.152 | 7.11 | 6.65 | 6.44 | 0.141 | 6.81 | 6.68 | 6.98 | 0.156 | 0.191 | 0.402 | 0.031 | 0.769 | 0.040 | 0.179 | 0.008 |
| MMP7 | multiparous | 4.56 | 3.48 | 4.37 | 0.563 | 3.26 | 3.63 | 3.69 | 0.558 | 4.94 | 4.21 | 3.88 | 0.576 | | | | | | | |
| | primiparous | 3.61 | 1.91 | 3.26 | 0.634 | 4.39 | 2.73 | 4.63 | 0.589 | 3.57 | 3.37 | 4.97 | 0.657 | 0.296 | 0.424 | 0.002 | 0.222 | 0.027 | 0.622 | 0.117 |
| MMP9 | multiparous | 5.80 | 5.93 | 6.07 | 0.240 | 5.76 | 5.68 | 6.12 | 0.237 | 5.98 | 6.14 | 6.12 | 0.246 | | | | | | | |
| | primiparous | 5.30 | 4.99 | 5.52 | 0.269 | 5.22 | 5.63 | 5.68 | 0.250 | 5.52 | 5.17 | 5.93 | 0.280 | 0.003 | 0.609 | 0.004 | 0.713 | 0.457 | 0.845 | 0.148 |

| | | | | | | | | | | | | | | | | | | | |
|---|---|---|---|---|---|---|---|---|---|---|---|---|---|---|---|---|---|---|---|
| MUC1 | multiparous | 5.03 | 5.09 | 5.21 | 0.228 | 5.10 | 4.93 | 4.73 | 0.226 | 5.36 | 5.33 | 4.92 | 0.233 | | | | | | | |
| | primiparous | 5.56 | 5.55 | 5.69 | 0.259 | 5.97 | 5.28 | 5.45 | 0.240 | 5.56 | 5.47 | 6.02 | 0.266 | 0.002 | 0.529 | 0.402 | 0.867 | 0.147 | 0.233 | 0.255 |
| NFKB1 | multiparous | 6.40 | 6.27 | 6.71 | 0.211 | 6.16 | 5.95 | 6.44 | 0.209 | 6.29 | 6.35 | 6.43 | 0.216 | | | | | | | |
| | primiparous | 6.13 | 5.80 | 6.28 | 0.236 | 6.11 | 5.91 | 6.20 | 0.220 | 6.24 | 5.88 | 6.60 | 0.247 | 0.167 | 0.592 | 0.001 | 0.691 | 0.491 | 0.994 | 0.417 |
| NLRP3 | multiparous | 7.19 | 7.08 | 7.54 | 0.154 | 7.01 | 7.06 | 7.30 | 0.152 | 7.28 | 7.21 | 7.26 | 0.158 | | | | | | | |
| | primiparous | 6.93 | 6.69 | 7.10 | 0.172 | 6.95 | 6.58 | 7.08 | 0.160 | 7.02 | 6.60 | 7.20 | 0.180 | 0.007 | 0.688 | < 0.001 | 0.904 | 0.051 | 0.833 | 0.370 |
| NFE2L2 | multiparous | 5.59 | 5.47 | 6.62 | 0.153 | 5.31 | 5.50 | 6.24 | 0.152 | 5.76 | 5.74 | 6.37 | 0.157 | | | | | | | |
| | primiparous | 5.31 | 5.19 | 6.66 | 0.173 | 4.96 | 5.35 | 6.48 | 0.161 | 5.21 | 5.12 | 6.60 | 0.179 | 0.063 | 0.292 | < 0.001 | 0.673 | 0.001 | 0.077 | 0.545 |
| PLAU | multiparous | 7.85 | 7.81 | 7.82 | 0.168 | 7.79 | 7.77 | 7.59 | 0.167 | 7.94 | 7.88 | 7.97 | 0.170 | | | | | | | |
| | primiparous | 7.71 | 7.39 | 7.83 | 0.193 | 7.72 | 7.44 | 7.23 | 0.178 | 7.79 | 7.33 | 7.72 | 0.195 | 0.003 | 0.165 | 0.148 | 0.836 | 0.273 | 0.310 | 0.875 |
| PTGS2 | multiparous | 7.57 | 7.65 | 5.88 | 0.201 | 7.45 | 7.49 | 6.13 | 0.200 | 7.36 | 7.54 | 6.24 | 0.203 | | | | | | | |
| | primiparous | 7.26 | 7.05 | 5.57 | 0.231 | 7.28 | 7.08 | 5.81 | 0.213 | 7.44 | 7.14 | 5.28 | 0.233 | 0.001 | 0.922 | < 0.001 | 0.854 | 0.219 | 0.795 | 0.413 |
| RIPK1 | multiparous | 5.55 | 5.56 | 5.93 | 0.153 | 5.41 | 5.46 | 5.56 | 0.152 | 5.49 | 5.59 | 5.68 | 0.155 | | | | | | | |
| | primiparous | 5.40 | 5.31 | 5.68 | 0.174 | 5.59 | 5.45 | 5.69 | 0.161 | 5.44 | 5.42 | 5.98 | 0.178 | 0.752 | 0.808 | 0.002 | 0.335 | 0.449 | 0.720 | 0.730 |
| S100A7 | multiparous | 4.44 | 4.51 | 4.31 | 0.331 | 4.23 | 4.25 | 4.42 | 0.329 | 4.68 | 5.05 | 4.45 | 0.338 | | | | | | | |
| | primiparous | 4.98 | 3.96 | 4.42 | 0.378 | 4.46 | 3.99 | 4.32 | 0.350 | 4.87 | 3.91 | 4.72 | 0.386 | 0.690 | 0.404 | 0.165 | 0.879 | 0.017 | 0.934 | 0.663 |
| S100A8 | multiparous | 7.58 | 7.56 | 7.55 | 0.143 | 7.52 | 7.56 | 7.58 | 0.142 | 7.65 | 7.69 | 7.63 | 0.147 | | | | | | | |
| | primiparous | 7.42 | 7.01 | 7.38 | 0.160 | 7.21 | 7.05 | 7.30 | 0.149 | 7.49 | 7.01 | 7.37 | 0.167 | 0.001 | 0.707 | 0.016 | 0.946 | 0.006 | 0.792 | 0.868 |
| S100A9 | multiparous | 7.61 | 7.55 | 7.51 | 0.150 | 7.54 | 7.53 | 7.58 | 0.148 | 7.67 | 7.69 | 7.57 | 0.154 | | | | | | | |
| | primiparous | 7.44 | 7.03 | 7.37 | 0.168 | 7.23 | 7.14 | 7.24 | 0.156 | 7.54 | 7.02 | 7.31 | 0.175 | 0.003 | 0.784 | 0.027 | 0.948 | 0.029 | 0.637 | 0.581 |
| SCD | multiparous | 7.00 | 7.15 | 6.74 | 0.186 | 7.13 | 6.94 | 6.69 | 0.185 | 7.23 | 7.16 | 6.67 | 0.190 | | | | | | | |
| | primiparous | 7.40 | 7.43 | 7.19 | 0.211 | 7.73 | 7.28 | 7.10 | 0.196 | 7.33 | 7.45 | 7.44 | 0.217 | 0.001 | 0.875 | 0.002 | 0.960 | 0.409 | 0.380 | 0.442 |
| SIRT1 | multiparous | 6.20 | 6.25 | 6.56 | 0.155 | 6.16 | 6.18 | 6.42 | 0.154 | 6.11 | 5.90 | 6.27 | 0.157 | | | | | | | |
| | primiparous | 6.05 | 6.17 | 6.65 | 0.177 | 5.71 | 6.39 | 6.41 | 0.164 | 5.77 | 6.17 | 6.67 | 0.180 | 0.946 | 0.335 | < 0.001 | 0.664 | 0.013 | 0.542 | 0.537 |
| SOCS3 | multiparous | 5.90 | 5.93 | 6.30 | 0.182 | 5.54 | 5.66 | 6.00 | 0.180 | 5.70 | 5.80 | 6.15 | 0.186 | | | | | | | |
| | primiparous | 5.33 | 5.40 | 5.62 | 0.205 | 5.29 | 5.51 | 5.80 | 0.191 | 5.45 | 5.37 | 5.77 | 0.213 | 0.003 | 0.728 | 0.001 | 0.409 | 0.927 | 0.919 | 0.964 |
| SOD1 | multiparous | 7.59 | 7.40 | 7.75 | 0.091 | 7.54 | 7.36 | 7.54 | 0.091 | 7.72 | 7.58 | 7.54 | 0.093 | | | | | | | |
| | primiparous | 7.81 | 7.68 | 7.96 | 0.105 | 7.92 | 7.59 | 7.80 | 0.097 | 7.67 | 7.74 | 8.05 | 0.106 | 0.001 | 0.256 | 0.001 | 0.767 | 0.410 | 0.250 | 0.104 |
| SOD2 | multiparous | 7.63 | 7.61 | 7.76 | 0.135 | 7.47 | 7.44 | 7.55 | 0.134 | 7.67 | 7.66 | 7.52 | 0.139 | | | | | | | |
| | primiparous | 7.42 | 7.11 | 7.41 | 0.15 | 7.52 | 7.29 | 7.33 | 0.14 | 7.43 | 7.14 | 7.52 | 0.158 | 0.021 | 0.866 | 0.006 | 0.593 | 0.032 | 0.762 | 0.170 |
| SPARC | multiparous | 4.15 | 4.04 | 3.85 | 0.174 | 4.19 | 4.29 | 3.88 | 0.173 | 3.93 | 4.44 | 4.09 | 0.176 | | | | | | | |
| | primiparous | 3.90 | 3.72 | 4.08 | 0.200 | 3.56 | 4.00 | 4.26 | 0.185 | 3.87 | 3.94 | 4.11 | 0.202 | 0.070 | 0.599 | 0.366 | 0.949 | 0.011 | 0.452 | 0.441 |
| STAT1 | multiparous | 7.09 | 6.84 | 7.21 | 0.178 | 6.93 | 6.86 | 7.03 | 0.176 | 7.12 | 6.82 | 6.98 | 0.183 | | | | | | | |
| | primiparous | 6.92 | 6.69 | 6.98 | 0.198 | 6.93 | 6.55 | 6.93 | 0.185 | 6.89 | 6.65 | 7.10 | 0.208 | 0.291 | 0.853 | 0.001 | 0.965 | 0.612 | 0.996 | 0.554 |
| STAT2 | multiparous | 4.18 | 4.11 | 4.36 | 0.159 | 4.08 | 4.01 | 4.21 | 0.157 | 4.03 | 4.14 | 4.29 | 0.162 | | | | | | | |
| | primiparous | 3.80 | 3.99 | 4.09 | 0.180 | 4.15 | 3.91 | 4.27 | 0.167 | 3.91 | 3.86 | 4.12 | 0.185 | 0.160 | 0.932 | 0.010 | 0.517 | 0.961 | 0.796 | 0.779 |
| STAT3 | multiparous | 6.97 | 7.00 | 7.24 | 0.129 | 6.85 | 6.95 | 7.03 | 0.128 | 6.94 | 6.92 | 6.98 | 0.132 | | | | | | | |
| | primiparous | 6.70 | 6.75 | 6.97 | 0.144 | 6.85 | 6.75 | 6.86 | 0.134 | 6.82 | 6.70 | 7.02 | 0.151 | 0.076 | 0.867 | 0.003 | 0.723 | 0.622 | 0.639 | 0.710 |
| STAT4 | multiparous | 6.28 | 6.18 | 6.35 | 0.168 | 6.33 | 5.91 | 6.05 | 0.166 | 6.43 | 6.17 | 6.29 | 0.171 | | | | | | | |
| | primiparous | 6.69 | 6.55 | 6.82 | 0.190 | 6.67 | 6.27 | 6.44 | 0.176 | 6.31 | 6.61 | 6.99 | 0.195 | 0.002 | 0.219 | 0.032 | 0.960 | 0.187 | 0.119 | 0.324 |
| STAT5A | multiparous | 6.09 | 6.00 | 6.49 | 0.165 | 6.01 | 5.92 | 6.16 | 0.164 | 6.13 | 6.12 | 6.23 | 0.168 | | | | | | | |
| | primiparous | 5.99 | 5.90 | 6.18 | 0.189 | 6.12 | 5.89 | 5.65 | 0.175 | 5.98 | 5.89 | 6.46 | 0.192 | 0.180 | 0.236 | 0.033 | 0.872 | 0.728 | 0.207 | 0.190 |
| TAP1 | multiparous | 6.23 | 5.81 | 6.13 | 0.326 | 5.99 | 5.73 | 5.49 | 0.323 | 6.70 | 6.51 | 6.39 | 0.334 | | | | | | | |
| | primiparous | 6.30 | 5.55 | 5.87 | 0.365 | 6.41 | 5.89 | 6.04 | 0.339 | 6.62 | 6.10 | 6.12 | 0.381 | 0.971 | 0.199 | 0.005 | 0.474 | 0.553 | 0.810 | 0.969 |
| TLR2 | multiparous | 6.70 | 6.60 | 6.54 | 0.153 | 6.49 | 6.69 | 6.47 | 0.151 | 6.76 | 6.74 | 6.40 | 0.157 | | | | | | | |
| | primiparous | 6.42 | 6.12 | 6.27 | 0.170 | 6.32 | 6.31 | 6.22 | 0.159 | 6.58 | 6.22 | 6.49 | 0.179 | 0.017 | 0.674 | 0.064 | 0.891 | 0.030 | 0.269 | 0.567 |
| TLR4 | multiparous | 6.90 | 6.87 | 6.95 | 0.149 | 6.73 | 6.92 | 6.84 | 0.147 | 6.86 | 6.97 | 6.69 | 0.152 | | | | | | | |
| | primiparous | 6.59 | 6.39 | 6.76 | 0.168 | 6.27 | 6.48 | 6.58 | 0.156 | 6.68 | 6.43 | 6.81 | 0.173 | 0.003 | 0.578 | 0.291 | 0.761 | 0.031 | 0.287 | 0.666 |
| TJP1 | multiparous | 5.29 | 5.64 | 4.02 | 0.186 | 5.40 | 5.28 | 4.27 | 0.185 | 5.42 | 5.39 | 4.52 | 0.188 | | | | | | | |
| | primiparous | 5.31 | 5.17 | 3.95 | 0.213 | 5.72 | 5.73 | 4.16 | 0.197 | 5.31 | 5.37 | 3.64 | 0.216 | 0.331 | 0.218 | < 0.001 | 0.071 | 0.126 | 0.975 | 0.184 |
| TNFA | multiparous | 7.27 | 7.32 | 5.22 | 0.209 | 7.17 | 7.28 | 5.60 | 0.208 | 7.05 | 7.23 | 5.54 | 0.212 | | | | | | | |
| | primiparous | 6.89 | 6.86 | 4.90 | 0.240 | 6.98 | 6.65 | 5.15 | 0.222 | 7.17 | 6.93 | 4.61 | 0.243 | 0.001 | 0.864 | < 0.001 | 0.977 | 0.237 | 0.572 | 0.428 |
| TXNRD1 | multiparous | 7.19 | 7.24 | 5.03 | 0.209 | 7.04 | 7.15 | 5.40 | 0.208 | 6.89 | 7.12 | 5.35 | 0.212 | | | | | | | |
| | primiparous | 6.82 | 6.84 | 4.69 | 0.241 | 6.95 | 6.92 | 4.92 | 0.223 | 6.97 | 6.83 | 4.42 | 0.243 | 0.001 | 0.558 | < 0.001 | 0.893 | 0.199 | 0.837 | 0.583 |
| reference genes 3 | multiparous | 10.73 | 10.61 | 10.63 | 0.184 | 10.40 | 10.65 | 10.55 | 0.183 | 10.55 | 10.58 | 10.25 | 0.188 | | | | | | | |
| | primiparous | 10.74 | 10.46 | 10.54 | 0.209 | 10.78 | 10.60 | 10.38 | 0.194 | 11.00 | 10.72 | 10.95 | 0.215 | 0.244 | 0.732 | 0.277 | 0.235 | 0.285 | 0.850 | 0.446 |

[1] Milk samples were collected on weeks 4, 8 et 12 after calving, then MEC from milk were prepared and an analysis of the gene expression antioxidant, immune and milk production function was made using ACTB (actin beta), GAPDH (Glyceraldehyde-3-Phosphate Dehydrogenase), PPIA (cyclophilin A), and RPLP0 (Ribosomal Protein Lateral Stalk Subunit P0) as reference genes.

[2] Probability of parity effects: TREAT = effect of treatment, control vs. vitamin E vs. plant extracts; PAR = effect of parity, primiparous vs. multiparous; WK = effect of week in week 4, 8 and 12 relative to calving; TREAT × WK = interaction between treatment and week; PAR × WK = interaction between parity and week; TREAT × PAR = interaction between treatment and parity; TREAT × PAR × WK = interaction between treatment and parity.

[3] Decimal logarithm of the geometric mean of ACTB (actin beta), GAPDH (Glyceraldehyde-3-Phosphate Dehydrogenase), PPIA (cyclophilin A), and RPLP0 (Ribosomal Protein Lateral Stalk Subunit P0) genes.

**Table S6**: Composition on dry matter ingredients and the chemical composition of the diet during close-up, first 15 (days in milk) DIM and after 15 DIM in dairy cows.

| Item [1] | Close-up [2] | Lactation first 15 DIM | Lactation after 15 |
|---|---|---|---|
| **Dry matter ingredients** | | | |
| Corn silage (% DM) | 72 | 65 | 65 |
| Energy concentrate (% DM) | 14 | 12.5 | 12.5 |
| Soybean meal (% DM) | 14 | 12.5 | 12.5 |
| Dehydrated alfalfa (% DM) | 0 | 10 | 10 |
| Straw (kg DM/day) | 1 | 1 | 0 |
| Mineral feed dry (g DM/day) [3] | 0.15 | 0 | 0 |
| Mineral feed lactation (g DM/day) [4] | 0 | 0.2 | 0.2 |
| Magnesium chloride (g DM/day) | 0.1 | 0 | 0 |
| Magnesium oxide (g DM/day) | 0 | 0.05 | 0.05 |
| Carbonates (g/kg DM) | 0 | 0.05 | 0.05 |
| **Chemical composition of the diet** | | | |
| DMI (kg/day) | 11.63 | 15.69 | 21.69 |
| $NE_L$ (MJ/kg DMI) [5] | 7.06 | 6.31 | 6.42 |
| $NE_L$ (MJ/day) [5] | 82.07 | 98.95 | 139.27 |
| DPI (g/kg DM) [6] | 82.27 | 79.97 | 84.16 |
| DPI (g/day) [6] | 956.57 | 1254.87 | 1825.77 |
| DPI/FU (g/FU) [6] | 85.95 | 93.51 | 96.67 |
| Nitrogen efficiency (g/g) | 0.47 | 0.95 | 0.8 |
| Calcium (g/kg DMI) | 4.44 | 9.13 | 7.93 |
| Phosphorus (g/kg DMI) | 3.33 | 3.54 | 3.37 |
| Magnesium (g/kg DMI) | 3.26 | 2.12 | 1.94 |
| DCAD (mEq/kg DMI) [7] | 187 | 234.16 | 237.73 |
| Selenium (g/kg DMI) | 0.31 | 0.47 | 0.36 |
| Vitamin E (IU/day) | 1257.97 | 994.21 | 1256.08 |

[1] With this diet composition, cows in the vitamin E group were supplemented with the all-rac-alpha-tocopheryl acetate form of vitamin E at a content of 30,000 IU/kg of this preparation (knowing that 1 mg equals 1 IU for all-rac-alpha-tocopheryl), cows received 3000 IU/day, then after calving for 12 weeks of lactation 1000 IU/day. For the plant extract group, the plant extracts were composed of *Sambucus nigra, Salix alba, Laurus nobilis, Haragophytum procumbens, Silybum marianum, Arctium lappa*, 0.06 mg/kg selenium and 1.56 µg/g dry matter gamma-tocopherol and 7.01 µg/g dry matter alpha-tocopherol.

[2] Before the close-up period (far-off period), the dry matter ingredients were corn silage (6.46% DM), grass silage (3.5% DM), rapeseed meal (1.32% DM), straw (0.88 kg DM/day) and mineral feed (150 g DM/day).

[3] The mineral feed lactation composition was the same before and after calving and was composed of calcium carbonate, magnesium sulfate, magnesium phosphate, wheat grinding, monocalcium phosphate containing a vitamin and trace mineral mixture supplying 250,000 IU/kg vitamin A, 60,000 IU/kg vitamin D3, 4,000 mg/kg vitamin E, 410 mg/kg Cu sulfate, 1,500 mg/kg Zn sulfate, 1,350 mg/kg Mn sulfate, 30 mg/kg I, 12 mg/kg Co, and 15 mg/kg Se.

[4] The mineral feed lactation composition was the same before and after calving and was composed of calcium carbonate, magnesium sulfate, magnesium phosphate, wheat grinding, monocalcium phosphate containing a vitamin and trace mineral mixture supplying 500,000 IU/kg vitamin A, 100,000 IU/kg vitamin D3, 1,500 mg/kg vitamin E, 960 mg/kg Cu sulfate, 3,630 mg/kg Zn oxide, 2,800 mg/kg Mn oxide, 80 mg/kg I, 35 mg/kg Co, and 22 mg/kg Se.

[5] NEL: Net energy for milk production in MJ/kg of DM or MJ/day.

[6] DPI: Digestible proteins in the intestine in g/kg of DM, or g/day, or g/FU (fodder unit) (INRA, 2018).

[7] DCAD: Dietary cation anion difference, calculated as the sum of the dietary contents in K and Na minus the dietary contents in Cl and S, with contents expressed in mEq/kg of DMI.

**Table S7**: Forward and reverse primers used for real-time quantitative PCR in mammary epithelial cells (MEC) to measure gene expression of antioxidant, immune and milk production functions.

| Item | Forward/Reverse | Primer sequence | Ref NCBI |
|---|---|---|---|
| ABCG2 | Forward | GGAGTCATGAAACCTGGCC | NM_001037478 |
| | Reverse | GGATCCTTCCTTGCAGCTAA | |
| AHR | Forward | GTGCAGAAAACTGTCAAGCCA | NM_001206026 |
| | Reverse | AACATCTGGTGGGAAAGGCAG | |
| AOAH | Forward | AACGACAGCAACAAGATGGC | NM_001078096.2 |
| | Reverse | AACACCCCAAATGCCGTTAC | |
| BAX | Forward | AGAAGCTGAGCGAGTGTCTGAA | NM_173894 |
| | Reverse | CGCTCTCGAAGGAAGTCCAA | |
| CASP1 | Forward | CTCCCACCTGGCAGGAATAC | XM_002692921 |
| | Reverse | AGGAGCTGGAAAAGGAGGGA | |
| CASP1P3 | Forward | TCCGGACATTCAACAACCGT | NM_176638.5 |
| | Reverse | ACCCACAATTCCCCACGATT | |
| CASP8 | Forward | AATATTGGGGAGCAGCTGGG | NM_001045970.2 |
| | Reverse | AGGCATCCTTGATGGGTTCC | |
| CAT | Forward | AGATGGACACAGGCACATGA | NM_001035386.2 |
| | Reverse | ACTGCCTCTCCATTTGCATT | |
| CCL2 | Forward | GCTCGCTCAGCCAGATGCAA | NM_174006 |
| | Reverse | GGACACTTGCTGCTGGTGACTC | |
| CCL20 | Forward | TTCGACTGCTGTCTCCGATA | NM_174263 |
| | Reverse | GCACAACTTGTTTCACCCACT | |
| CCL5 | Forward | CTGCCTTCGCTGTCCTCCTGATG | NM_175827 |
| | Reverse | TTCTCTGGGTTGGCGCACACCTG | |
| CD14 | Forward | TCCACAGTCCAGCCGACAAC | NM_174008 |
| | Reverse | AACGGCGCTAGACCAGTCAG | |
| CD36 | Forward | CTGGCTGTGTTTGGAGGGATTC | NM_001278621 |
| | Reverse | ACTGTCACTTCATCTGGATTCTGC | |
| CDH1 | Forward | TGCCCGACCATCAAAGAGAT | NM_174031 |
| | Reverse | CGGTACGAAATGGCTTCCAC | |
| CLDN1 | Forward | GAAGAACAGCACGTACACGGC | NM_001002763 |
| | Reverse | CCTCTGGCAGAAGTCCATGGT | |
| CLEC4A | Forward | AAGACGACGAGGCACAGAAG | NM_001001854 |
| | Reverse | AGCCCAGCCAATGAAGAGA | |
| CLEC7A | Forward | GAAGTTACCACCGTGCTTGC | NM_001191510.1 |
| | Reverse | CCTCTGAAGTCATGCTGCGA | |
| CSN1S1 | Forward | AGGCAAGTGTCTTCCCAGC | NM_001031852.1 |
| | Reverse | ACAACAAGGTGGAGCCATCC | |
| CSN3 | Forward | GTTCCCAATAGTGCTGAGGAACGACT | NM_181029.2 |
| | Reverse | GGGTAGAAGTAGGCCAGTTCCTGAT | |
| CTSB | Forward | TGCAATGATGAAGAGTTTTTTCCTAG | NM_174294 |
| | Reverse | GATTGGGATATATTTGGCTATTTTGT | |
| CXCL10 | Forward | TTCAGGCAGTCTGAGCCTAC | NM_001046551 |
| | Reverse | ACGTGGGCAGGATTGACTTG | |
| CXCL2 | Forward | GTGTCTCAACCCCGCCGCTC | NM_174299.3 |
| | Reverse | TCCAGATGGCCTTAGGAGGTGG | |
| CXCL8 | Forward | TGAAGCTGCAGTTCTGTCAAG | NM_173925.2 |

| | | | |
|---|---|---|---|
| | Reverse | TTCTGCACCCACTTTTCCTTGG | |
| DEFB5 | Forward | TCGTGCTCCTCTTCCTAGTC | NM_001130761 |
| | Reverse | GGCACGAGATCGGAATACAG | |
| RIG-I | Forward | TGTGGTGGAAGATGTTGCGA | XM_002689480 |
| | Reverse | AGGGGACATTTCTGCAGCAT | |
| FABP3 | Forward | GCGTTCTCTGTCGTCTTTCCC | NM_174313 |
| | Reverse | CTGTGTTCTTGAAGGTGCTTTGTG | |
| FAS | Forward | GCCCACATGGCTGGTATCAA | NM_174662.2 |
| | Reverse | TTTTTCCGTTTGCCAGGAGG | |
| GPX1 | Forward | GCAAGGTGCTGCTCATTGAG | NM_174076 |
| | Reverse | CGCTGCAGGTCATTCATCTG | |
| GPX3 | Forward | GTCAACGTGGCCAGCTACTGA | NM_174077 |
| | Reverse | CAGAATGACCAGACCAAATGGTT | |
| HSPA8 | Forward | CGAATCATCAATGAGCCAACTG | NM_174345 |
| | Reverse | TGCCACCCCCTAAATCAAAG | |
| IFNAR1 | Forward | TCCTTTGCCACGTGTCAAGT | NM_174552.2 |
| | Reverse | AGTAGCGTGAGGGAGACAGA | |
| IFNAR2 | Forward | GTGTGGGTAAACACGACGGA | NM_174553 |
| | Reverse | TGGGTCCAAAGGCTTGTCTG | |
| IFNB1 | Forward | GACAGTCGTGCAAGTGCAAA | NM_174350.1 |
| | Reverse | CAGTCACGGACGTAACCTGT | |
| IL10 | Forward | GTGATGCCACAGGCTGAGAA | NM_174088.1 |
| | Reverse | TGCTCTTGTTTTCGCAGGGCAG | |
| IL1A | Forward | CTGAAGAAGAGACGGTTGAG | NM_174092 |
| | Reverse | ATGCATTCCTGGTGGATGAC | |
| IL1B | Forward | CTCTCACAGGAAATGAACCGAG | EU276067 |
| | Reverse | GCTGCAGGGTGGGCGTATCACC | |
| IL2RA | Forward | AACACACAGATGCGCAGAAC | NM_174358 |
| | Reverse | TTCACGTTCGTGTTCCCATG | |
| IL33 | Forward | GATGGTGGCAGTCATCGGAA | NM_001075297.1 |
| | Reverse | GTAGCTCCACAGAGTGCTCC | |
| IL6 | Forward | TGCTGGTCTTCTGGAGTATC | EU276071 |
| | Reverse | GTGGCTGGAGTGGTTATTAG | |
| IRF3 | Forward | GGAAGGATAAGCCCGACCTG | NM_001029845.3 |
| | Reverse | GAGTCCTTGCTGTGGTCCTC | |
| ITGA4 | Forward | GGGTTTGTAAGTCCAGCCTCA | NM_174748.1 |
| | Reverse | CGGTGTTGATGACGTGGAAAG | |
| LALBA | Forward | ACCAGTGGTTATGACACACAAGC | NM_174378 |
| | Reverse | AGTGCTTTATGGGCCAACCAGT | |
| LAP | Forward | TGCTCCTTGCGCTCCTCTTC | NM_203435 |
| | Reverse | CTCCGAGACAGGTGCCAATC | |
| LBP | Forward | CCTGATTCTAGCATTCGACAG | NM_001038674 |
| | Reverse | GCTGAAGTTCAGGCACG | |
| LCN2 | Forward | TGGCGGGGAATGCAATTAAG | XM_002691670 |
| | Reverse | TAACAGGATGGAGGTGACGTTG | |
| LPL | Forward | AGTTTATGAACTGGATGGCGGATG | NM_001075120 |
| | Reverse | TCAGGAGAAAGGCGACTTGGAG | |
| MMP7 | Forward | GGAGCGAAGCAATCCCACTG | NM_001075130 |

| | | | |
|---|---|---|---|
| | Reverse | GGCCCATCAAAGGGATATGGG | |
| MMP9 | Forward | CGTTCCGACGACATGCTCTG | NM_174744 |
| | Reverse | CATTGCCGTCCTGGGTGTAG | |
| MUC1 | Forward | CTCTCCAGGCCATGATAGTG | NM_174115 |
| | Reverse | AAGTGACCATGGAGCTTGAC | |
| NFE2L2 | Forward | AGGACATGGATTTGATTGAC | NM_001011678 |
| | Reverse | TACCTGGGAGATGTTGGCA | |
| NFKB1 | Forward | TGAGGCCATTGACGTGATCC | NM_001076409 |
| | Reverse | TGCGGAAGGAGGTCTCTACA | |
| NLRP3 | Forward | CTCAGTGGCAATACCCTGGG | NM_001102219.1 |
| | Reverse | AGCACTGTCCCAACCACAAT | |
| PLAU | Forward | GCCAGGAGTCTACACAAGGG | NM_174147 |
| | Reverse | TGGGGTCCTTCAGAGGACAA | |
| PTGS2 | Forward | CATGGGTGTGAAAGGGAGGAA | AF004944 |
| | Reverse | ATTTGTGCCCTGGGGATCAG | |
| RIPK1 | Forward | GCGAATCTCTCGGGTTGTGT | NM_001035012 |
| | Reverse | GCCTGCTCCAGGAAGTCTG | |
| S100A7 | Forward | CAGCTTGAGCAGGCCATTAC | NM_174596 |
| | Reverse | CGTGGCTGTGGTTGTGATAG | |
| S100A8 | Forward | CTCCCTGATTGACGTCTACC | NM_001113725 |
| | Reverse | TCCAGGCCCACCTTTATCAC | |
| S100A9 | Forward | TGACACCCTGATCCAGAAAG | NM_001046328 |
| | Reverse | GCCACCAGCATAATGAACTC | |
| SCD | Forward | GCCTCTGCGGGTCTTCCTG | NM_173959 |
| | Reverse | GGTGGGCACGGTGATCTCG | |
| SIRT1 | Forward | ATACACTGGAGCAGGTT | NM_001192980 |
| | Reverse | TTCATCAGCTGGGCATCTAG | |
| SOCS3 | Forward | GATCCCTCTGGTGTTGAGCC | NM_174466.2 |
| | Reverse | CAGCTGGGTGACTTTCTCGT | |
| SOD1 | Forward | TGTTGCCATCGTGGATATTGTAG | NM_174615 |
| | Reverse | CCCAAGTCATCTGGTTTTTCATG | |
| SOD2 | Forward | GCAATTCCCCTTGGGGTTCT | NM_201527.2 |
| | Reverse | CAGCCTGCGTTGAAGTTCAA | |
| SPARC | Forward | GAGAATTCGATGATGGTGC | NM_174464 |
| | Reverse | GAAGGTCTTGTTGTCGTTG | |
| STAT1 | Forward | CAAAGGAAGCCCCAGAGCCTAT | NM_001077900.1 |
| | Reverse | GCCACTCTTCTGTGTTCACTTAC | |
| STAT2 | Forward | AGCCCGTTTCAGGATCAGC | NM_001205689.1 |
| | Reverse | CAGTGCAGCTTTCTGCCAGTTC | |
| STAT3 | Forward | GACCGGTGTCCAGTTCACAA | NM_001012671 |
| | Reverse | AAATTTCCGGGACCCTCTGA | |
| STAT4 | Forward | ACAGCAAATCGCCTGCATTG | NM_001083692 |
| | Reverse | CTCCAACTGCCGTCTGAGTT | |
| STAT5A | Forward | CATGTACCCACAGAACCCTGAC | NM_001012673.1 |
| | Reverse | GGGAGAGAGGGCTCCAGACT | |
| TAP | Forward | GTAGGAAATCCTGTAAGCTGTG | AF014106 |
| | Reverse | GTGTCTTGGCCTTCTTTTAC | |
| TJP1 | Forward | AATGCATCCTGACCACCAGG | XM_024982002 |

| | | | |
|---|---|---|---|
| | Reverse | GATGGTGCCGGGTTTGTTTC | |
| TLR2 | Forward | ACTGGGTGGAGAACCTCATGGTCC | NM_174197.2 |
| | Reverse | ATCTTCCGCAGCTTACAGAAGC | |
| TLR4 | Forward | GCATGGAGCTGAATCTCTAC | NM_174198.6 |
| | Reverse | CAGGCTAAACTCTGGATAGG | |
| TNF | Forward | TCTTCTCAAGCCTCAAGTAACAAGC | NM_173966.3 |
| | Reverse | CCATGAGGGCATTGGCATAC | |
| TXNRD1 | Forward | GACCAAACCATCGAAGGAGAGTAT | NM_174625 |
| | Reverse | CTCGTGCAAGCATCTCTTCCT | |
| ACTB | Forward | ACGGGCAGGTCATCACCATC | BT030480 |
| | Reverse | AGCACCGTGTTGGCGTAGAG | |
| GAPDH | Forward | GGCATCGTGGAGGGACTTATG | DQ403066 |
| | Reverse | GCCAGTGAGCTTCCCGTTGAG | |
| PPIA | Forward | TCCGGGATTTATGTGCCAGGG | BC105173 |
| | Reverse | GCTTGCCATCCAACCACTCAG | |
| RPLP0 | Forward | CAACCCTGAAGTGCTTGACAT | NM_001012682 |
| | Reverse | AGGCAGATGGATCAGCCA | |
